# Missense Mutations in the Human Nanophthalmos Gene *TMEM98* Cause Retinal Defects in the Mouse

**DOI:** 10.1101/513846

**Authors:** Sally H. Cross, Lisa Mckie, Margaret Keighren, Katrine West, Caroline Thaung, Tracey Davey, Dinesh C. Soares, Luis Sanchez-Pulido, Ian J. Jackson

## Abstract

**PURPOSE:** We previously found a dominant mutation, *Rwhs*, causing white spots on the retina accompanied by retinal folds. Here we identify the mutant gene to be *Tmem98.* In humans, mutations in the orthologous gene cause nanophthalmos. We modelled these mutations in mice and characterised the mutant eye phenotypes of these and *Rwhs*.

**METHODS:** The *Rwhs* mutation was identified to be a missense mutation in *Tmem98* by genetic mapping and sequencing. The human *TMEM98* nanophthalmos missense mutations were made in the mouse gene by CRISPR-Cas9. Eyes were examined by indirect ophthalmoscopy and the retinas imaged using a retinal camera. Electroretinography was used to study retinal function. Histology, immunohistochemistry and electron microscopy techniques were used to study adult eyes.

**RESULTS:** An I135T mutation of *Tmem98* causes the dominant *Rwhs* phenotype and is perinatally lethal when homozygous. Two dominant missense mutations of *TMEM98*, A193P and H196P are associated with human nanophthalmos. In the mouse these mutations cause recessive retinal defects similar to the *Rwhs* phenotype, either alone or in combination with each other, but do not cause nanophthalmos. The retinal folds did not affect retinal function as assessed by electroretinography. Within the folds there was accumulation of disorganised outer segment material as demonstrated by immunohistochemistry and electron microscopy, and macrophages had infiltrated into these regions.

**CONCLUSIONS:** Mutations in the mouse orthologue of the human nanophthalmos gene *TMEM98* do not result in small eyes. Rather, there is localised disruption of the laminar structure of the photoreceptors.

## INTRODUCTION

There are a range of genetic disorders which present with a reduced eye size. In microphthalmia the reduced size is often associated with additional developmental eye defects, such as coloboma, and may also include developmental defects in other organs. In some cases there is an overall size reduction without other developmental defects. The smaller eye may be a result of a reduction in size of the anterior segment alone (anterior microphthalmos), the posterior segment alone (posterior microphthalmos) or a reduction of both. This last category includes simple microphthalmos and the more severe nanophthalmos ^1, 2^. Posterior microphthalmos is sometimes considered as part of a continuum with nanophthalmos, as they have pathological and genetic features in common. In nanophthalmos eye length is reduced by 30% or more, and is usually associated with other ocular features, notably a thickened choroid and sclera, as well as a high incidence of glaucoma, corneal defects, vascular defects and a range of retinal features including retinitis pigmentosa, retinoschisis, retinal detachments and retinal folds. Retinal folds are also observed in posterior microphthalmos, and are sometimes ascribed to a relative overgrowth of neural retina within a smaller globe, resulting in folding of the excess retinal tissue.

A number of genes have been found to be associated with nanophthalmos ^2^. Mutations in three genes have established associations with nanophthalmos. Several families with nanophthalmos have been found to have clear loss of function mutations in both alleles of the gene encoding membrane-type frizzled related protein, *MFRP*, that is expressed principally in the RPE and ciliary body ^3^. Homozygous loss of function mutations in *MFRP* have been found in other individuals with posterior microphthalmia plus retinitis pigmentosa, foveoschisis and drusen, indicating likely genetic background effects on the severity or range of the disease ^4–6^. A second autosomal recessive nanophthalmos gene is *PRSS56*, encoding a serine protease. Families with biallelic mutations in this gene have been characterised, some of whom have posterior microphthalmia whilst others have nanophthalmos ^7–9^. There is no apparent genotype-phenotype correlation; there are patients with homozygous frameshift mutations with either condition. Intriguingly, association of variants at *PRSS56* with myopia has been reported in genome-wide association studies ^10, 11^.

Most recently three families have been characterised in which heterozygous mutations in *TMEM98* are segregating with nanophthalmos. Two families have missense mutations, the third has a 34bp deletion that removes the last 28 bases of exon 4 and the first six bases of intron 4 including the splice donor site^12, 13^. The effect of the deletion on the *TMEM98* transcript is unknown but *in silico* splicing prediction programs (Alamut Visual Splicing Predictions) predict that it could result in exon 4 being skipped, and the production of a potentially functional protein with an internal deletion. Alternatively, other splice defects would result in frameshifts, nonsense mediated decay and loss of protein from this allele. In this case the cause of the disease would be haploinsufficiency. Caution in assigning a role for *TMEM98* in nanophthalmos has been raised by findings in another study where different heterozygous missense mutations in *TMEM98* were found in patients with high myopia and cone-rod dystrophy ^14^.

Patients with heterozygous or homozygous mutations in the *BEST1* gene can have a range of defects ^15^. *BEST1* encodes the bestrophin-1 protein, a transmembrane protein located in the basolateral membrane of the retinal pigment epithelium (RPE) ^16^. The predominant disease resulting from heterozygous mutations in *BEST1* is Best vitelliform macular dystrophy, in which subretinal lipofuscin deposits precede vision defects ^17^. These patients in early stages have a normal electroretinogram (ERG) but have a defect in electrooculography (EOG) indicative of an abnormality in the RPE. This can progress to retinitis pigmentosa. Five families have been reported with dominant vitreoretinochoroidopathy and nanophthalmos due to three different missense mutations in *BEST1*. Each mutant allele can produce two isoforms, one containing a missense mutation and one containing an in-frame deletion ^18^. On the other hand, homozygous null mutations of *BEST1* result in high hyperopia accompanied by abnormal ERGs and EOGs and RPE abnormalities but lacking the vitelliform lesions found in the dominant disease^19^. Similar associations have been seen for mutations in the *CRB1* gene that encodes an apical transmembrane protein important for determining cell polarity in photoreceptors ^20^. *CRB1* mutations are most frequently found associated with recessive retinitis pigmentosa or with Leber congenital amaurosis and the disease phenotype observed in patients is very variable suggestive of the influence of genetic modifiers ^21, 22^. In addition, hypermetropia and short axial length are a common associations^23^. Furthermore, in two cases, both involving consanguineous families, the retinal dystrophy is associated with nanophthalmos ^24, 25^.

Mouse models with mutations in most of these genes have been analysed. Mice which have targeted disruption of the *Best1* gene do not recapitulate the human bestrophinopathy phenotype or have only an enhanced response in EOG (indicative of a defect in the RPE) ^26, 27^. However, when the common human mutation, W93C, is engineered into the mouse gene, both heterozygous and homozygous mice show a distinctive retinal pathology of fluid or debris filled retinal detachments which progress with age ^28^. Spontaneous and targeted mutations in mouse *Crb1* have been characterised, and they show similar, though not identical, recessive retinal phenotypes that had variable ages of onset ^29–31^. The defects found are focal. Folds appear in the photoreceptor layer that are visualised as white patches on retinal imaging. In a complete loss-of-function allele and the frameshift allele *Crb1*^*rd8*^ the folds are adjacent to discontinuities in the outer (external) limiting membrane (OLM) accompanied by loss of adherens junctions, and within them the photoreceptors, separated from the RPE, show degeneration ^29, 30^. In mice engineered to carry a missense mutation, *Crb1*^*C249W*^, that causes retinitis pigmentosa in humans, although the OLM appears intact, retinal degeneration still occurs, albeit later than in the other models ^31^. Similar to the human phenotype the extent of the retinal spotting observed in *Crb1* mutants is strongly affected by the genetic background.

Two lines of mice with spontaneous mutations in *Mfrp* have been described ^32–34^. These do not recapitulate the nanophthalmic phenotype observed in humans. Instead both have the same recessive phenotype of white spots on the retina, which correlate with abnormal cells below the retina that stain with macrophage markers, and progress to photoreceptor degeneration. For the *Mfrp*^*rdx*^ mutation, atrophy of the RPE was reported ^33^ and for *Mfrp*^*rd6*^ a modest ocular axial length reduction from about 2.87 to 2.83 mm was reported although apparently not statistically significant ^35^. A screen for mice with increased intraocular pressure (IOP) found a splice mutation in the *Prss56* gene predicted to produce a truncated protein ^8^. The increased IOP was associated with a narrow or closed iridocorneal angle, analogous to that seen in human angle closure glaucoma. In addition these mice have eyes that are smaller than littermates, although the size reduction is variable and slight, ranging from 0 to 10% decrease in axial length. Reduction in axial length only becomes statistically significant after post-natal day seven. More recently it has been shown that mice deficient for *Prss56* have eyes with a decreased axial length and hyperopia ^36^.

In summary the human nanophthalmos mutations modelled in mice produce either a much less severe, or non-significant, axial length reduction (*Prss56*, *Mfrp*) or no effect on eye size (*Best1*, *Crb1*). There are differences in the development of human and mouse eyes which probably underlies the differences in mutant phenotype.

To date no mouse models of *TMEM98* have been reported. We describe here characterisation of a mouse mutation in *Tmem98*, which results in a dominant phenotype of retinal folds. In addition we engineer the two human nanophthalmos-associated missense mutations of *TMEM98* into the mouse gene and show that these mice also, when homozygous or when compound heterozygous, have the same retinal fold phenotype but do not have a statistically significant reduction in eye size.

## MATERIALS AND METHODS

### Mice

All mouse work was carried out in compliance with UK Home Office regulations under a UK Home Office project licence and experiments adhered to the ARVO Statement for the Use of Animals in Ophthalmic and Vision Research. Clinical examinations were performed as previously described ^37^. Fundus imaging was carried out as described ^38^. Mice carrying a targeted knockout-first conditional-ready allele of *Tmem98, Tmem98*^*tm1a(EUCOMM)Wtsi*^ (hereafter *Tmem98*^*tm1a*^), were obtained from the Sanger Institute ^39^. *Tmem98*^*tm1a*/+^ mice were crossed with mice expressing Cre in the germ-line to convert this ‘knockout-first’ allele to the reporter knock-out allele *Tmem98*^*tm1b(EUCOMM)Wtsi*^ (hereafter *Tmem98*^*tm1b*^). In this allele the DNA between the loxP sites in the targeting cassette which includes the neo selection gene and the critical exon 4 of *Tmem98* is deleted. To create the *Tmem98*^*H196P*^ allele the CRISPR design site http://www.crispr.mit.edu was used to design guides and the selected guide oligos ex7_Guide1 and ex7_Guide2 (Supplementary Table S1) were annealed and cloned into the *Bbs I* site of the SpCas9 and chimeric guide RNA expression plasmid px330 ^40^ (pX330-U6-Chimeric_BB-CBh-hSpCas9 was a gift from Feng Zhang (Addgene plasmid #42230, https://www.addgene.org/)). Following pronuclear injection of this plasmid along with repair oligo H196P (Supplementary Table S1) the injected eggs were cultured overnight to the 2-cell stage and transferred into pseudopregnant females. To create the *Tmem98*^*A193P*^ allele Edit-R crRNA (sequence 5’-CCAAUCACUGUCUGCCGCUG-3’) (Dharmacon) was annealed to tracrRNA (Sigma) in IDT Duplex buffer (IDT). This, along with Geneart Platinum Cas9 nuclease (Invitrogen B25641) and repair oligo A193P (Supplementary Table S1) were used for pronuclear injection as described above. Pups born were screened for the targeted changes by sequencing PCR fragments generated using the oligos ex7F and ex7R (Supplementary Table S2) and lines established carrying the targeted missense mutations. Genotyping was initially done by PCR, and sequencing where appropriate, using the primers in Supplementary Table S2. Subsequently most genotyping was performed by Transnetyx using custom designed assays (http://www.transnetyx.com). All lines were maintained on the C57BL/6J mouse strain background.

### DNA Sequencing

The candidate interval was captured using a custom Nimblegen array and sequenced with 454 technology by Edinburgh Genomics (formerly known as GenePool) (http://www.genomics.ed.ac.uk).

### Bioinformatics

Multiple protein sequence alignments were generated with the program Clustal Omega^41^. The possible impact of missense mutations on protein structure and function was evaluated using Polyphen-2 (http://genetics.bwh.harvard.edu/pph2/)^42^. HMM-HMM comparison was used for protein homology detection^43^. For protein structure prediction Phyre2 was used (http://www.sbg.bio.ic.ac.uk/phyre2/)^44^. We used PyMOL for protein structure visualization^45^. SCWRL^46^ and the FoldX web server^47^ were used to evaluate the effect of amino acid substitutions on protein stability.

### Electroretinography

Prior to electroretinography mice were dark adapted overnight (>16 hours) and experiments were carried out in a darkened room under red light using an HMsERG system (Ocuscience). Mice were anesthetised using isofluorane and their pupils dilated by the topical application of 1% w/v tropicamide. Three grounding electrodes were placed subcutaneously (tail, and each cheek) and silver embedded electrodes were positioned on the corneas using hypromellose eye drops (2.5% methylcellulose coupling agent) held in place with a contact lens. Animals were kept on a heated platform to maintain them at 37°C and monitored using a rectal thermometer. A modified QuickRetCheck (Ocuscience) protocol was used for obtaining full-field scotopic ERGs. Briefly, 4 flashes at 10 mcd.s/m^2^ at 2 s intervals were followed by 4 flashes at 3 cd.s/m^2^ (at 10 s intervals) and then 4 flashes at 10 cd.s/m^2^ (at 10 s intervals).

### Histology and Immunostaining

Mice were culled, eyes enucleated and placed into Davidson’s fixative (28.5% ethanol, 2.2% neutral buffered formalin, 11% glacial acetic acid) for 1 hour (cryosectioning) or overnight (wax embedding) except for the eye used for Fig. 1 which was placed in 10% neutral buffered formalin for 24 hours before immersion in Davidson’s fixative. Prior to wax embedding eyes were dehydrated through an ethanol series. Haematoxylin and eosin staining was performed on 5 or 10 µm paraffin embedded tissue sections and images captured using a Nanozoomer XR scanner (Hamamatsu) and viewed using NDP.view2 software. For cryosectioning, fixed eyes were transferred to 5% sucrose in PBS and once sunk transferred to 20% sucrose in PBS overnight. Eyes were then embedded in OCT compound and cryosectioned at 14 µM. For immunostaining on cryosections, slides were washed with water then PBS and post-fixed in acetone at −20°C for 10 minutes. They were then rinsed with water, blocked in 10% donkey serum (DS), 0.1% Tween-20 in TBS (TBST) for one hour and then incubated with primary antibodies diluted in TBST with 5% DS for two hours at room temperature or overnight at 4°C. Subsequently, after washing with TBST, the slides were incubated with Alexa Fluor secondary antibodies (Invitrogen) diluted 1:400 in TBST with 5% DS at room temperature for one hour. Following washing with TBST coverslips were mounted on slides in Prolong Gold (ThermoFisher Scientific) and confocal images acquired on a Nikon A1R microscope. Images were processed using either NIS-Elements or ImageJ software.

**Figure 1.**
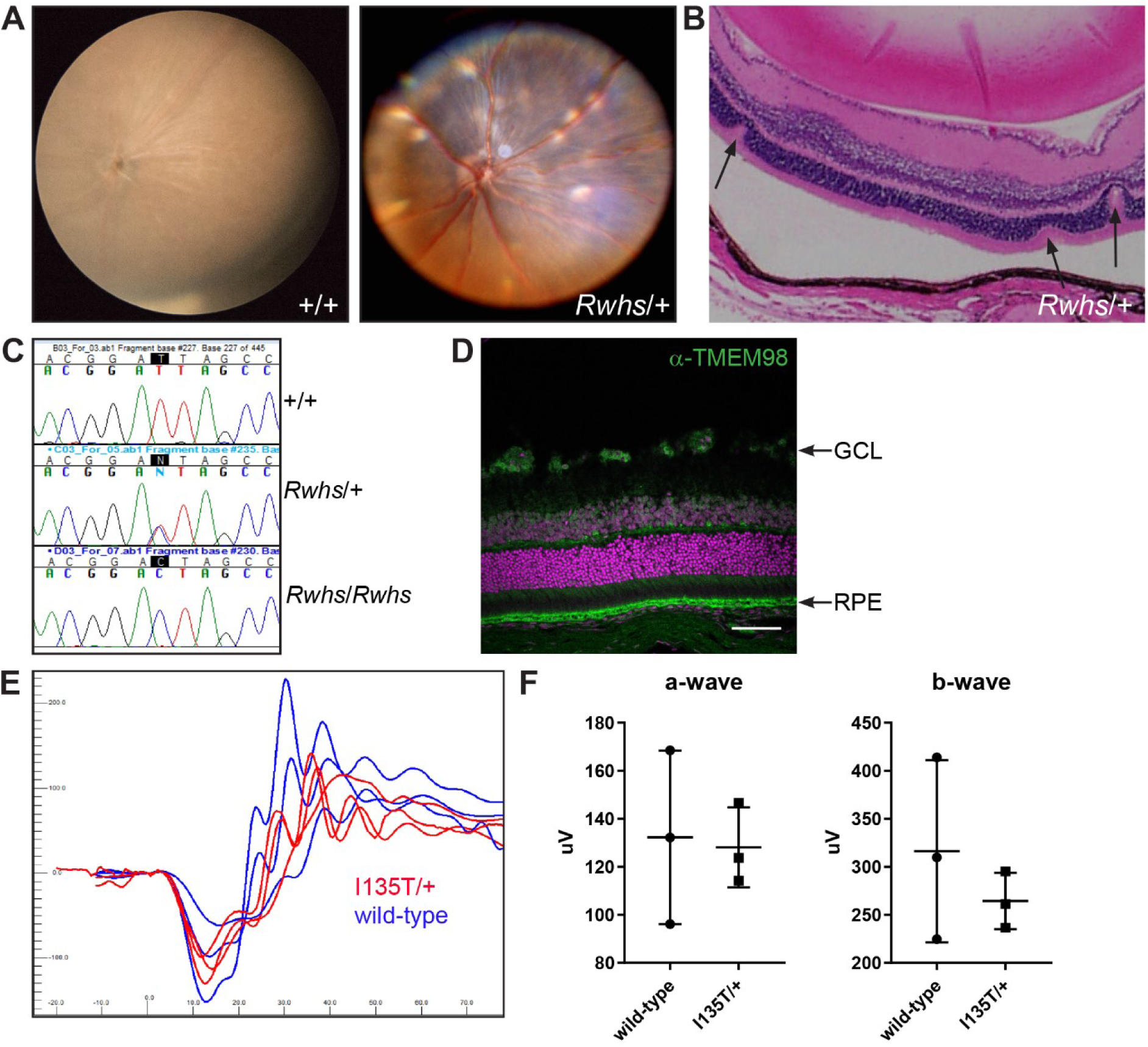
The *Rwhs* mutation is caused by an I135T mutation of the transmembrane gene *Tmem98*. (**A**) Retinal images of wild-type (+/+) and *Rwhs*/+ eyes. Scattered white spots are present on the *Rwhs*/+ retina. (**B**) *Rwhs*/+ retinal section with three folds in the outer nuclear layer indicated by arrows. The separation of the RPE from the outer segments is a fixation artefact (**C**) Genomic DNA sequence traces from exon 5 of *Tmem98* from wild-type (+/+), heterozygous mutant (*Rwhs*/+) and homozygous mutant (*Rwhs*/*Rwhs*) embryonic samples. The position of the T-to-C transition at position 404 (404T>C) in the coding sequence of *Tmem98* is highlighted. (**D**) A section of wild-type retina immunostained for TMEM98 (green). Prominent staining is seen in the retinal pigment epithelium (RPE) and there is also some staining in the ganglion cell layer (GCL). DNA is shown in magenta. (**E**) *Tmem98*^*I135T/+*^ mice have a normal ERG response. Shown are the responses at 3 cd.s/m^2^ (average of 4 flashes) for the left eye of three *Tmem98*^*I135T/+*^ mice (5-6 months of age) in red and three wild-type mice (5 months of age) in blue. (**F**) Comparison of a-wave amplitudes (right) and b-wave amplitudes (left), average of left and right eye for each mouse. There is no significant difference between *Tmem98*^*I135T/+*^ and wild-type mice (a-wave, unpaired t test with Welch’s correction, P = 0.87 and b-wave, unpaired t test with Welch’s correction, P = 0.45).Scale bar: 100 µM (**D**).

### Eye size measurment

Eyes from male mice were enucleated and kept in PBS prior to measuring to prevent drying out. The analysis was restricted to males as significant differences in axial length have been reported in the C57BL/6 strain between male and female mice^48^. Axial length measurements were made from the base of the optic nerve to the centre of the cornea using a Mitutoyo Digital ABS Caliper 6”/150mm, item number: 500-196-20.

### Antibodies

Primary antibodies used are listed in Table 1. DNA was stained with TOTO-3 (Invitrogen) or 4’,6-Diamidine-2’-phenylindole dihydrochloride (DAPI).

**Table 1.** Primary antibodies.

| Antibody | Source | Product No | Concentration used |
| --- | --- | --- | --- |
| anti-TMEM98 | proteintech | 14731-1-AP | 1:5000 (WB), 1:100 (IF) |
| anti-α-Tubulin | Sigma-Aldrich | T5168 | 1:10,000 (WB) |
| anti-Prominin 1 | proteintech | 18470-1-AP | 1:100 (IF) |
| anti-Rhodopsin | Millipore | MAB5356 | 1:500 (IF) |
| anti-ML Opsin | Millipore | AB5405 | 1:500 (IF) |
| anti-GFAP | abcam | ab7260 | 1:500 (IF) |
| anti-β-Catenein | Cell Signaling Technology | 19807S | 1:500 (IF) |
| Anti-F4/80 | AbD Serotec | MCA497EL | 1:500 (IF) |

Key: WB=Western blotting, IF=immunofluorescence

### Transmission Electron Microscopy

Samples were fixed in 2% EM grade glutaraldehyde (TAAB Laboratory Equipment, Aldermaston, UK) in sodium cacodylate buffer at 4°C overnight, post-fixed in 1% osmium tetroxide (Agar Scientific, Essex, UK), dehydrated in increasing concentrations of acetone and impregnated with increasing concentrations of epoxy resin (TAAB Laboratory Equipment). Embedding was carried out in 100% resin at 60°C for 24 hours. Semi-thin survey sections of 1 μm, stained with toluidine blue, were taken to determine relevant area. Ultrathin sections (approximately 70 nm) were then cut using a diamond knife on a Leica EM UC7 ultramicrotome (Leica, Allendale, NJ, USA). The sections were stretched with chloroform to eliminate compression and mounted on Pioloform-filmed copper grids (Gilder Grids, Grantham UK). To increase contrast the grids were stained with 2% aqueous uranyl acetate and lead citrate (Leica). The grids were examined using a Philips CM 100 Compustage (FEI) Transmission Electron Microscope. Digital images were collected using an AMT CCD camera (Deben UK Ltd., Suffolk, UK).

### Statistics

For the data shown in Figures 1, 2,3 and Supplementary Figure S7 the graphs were created and unpaired t tests with Welch’s correction were performed using the program Graphpad Prism. For the data shown in Tables 2, S3 and S4 chi square tests were performed using http://graphpad.com/quickcalcs. A value of P<0.05 was considered significant.

**Table 2.**
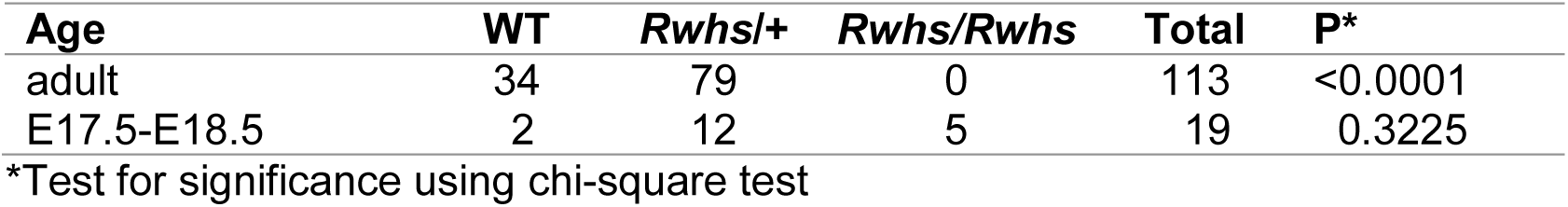
*Tmem98*^*Rwhs/+*^ intercross genotyping results.

**Figure 2.**
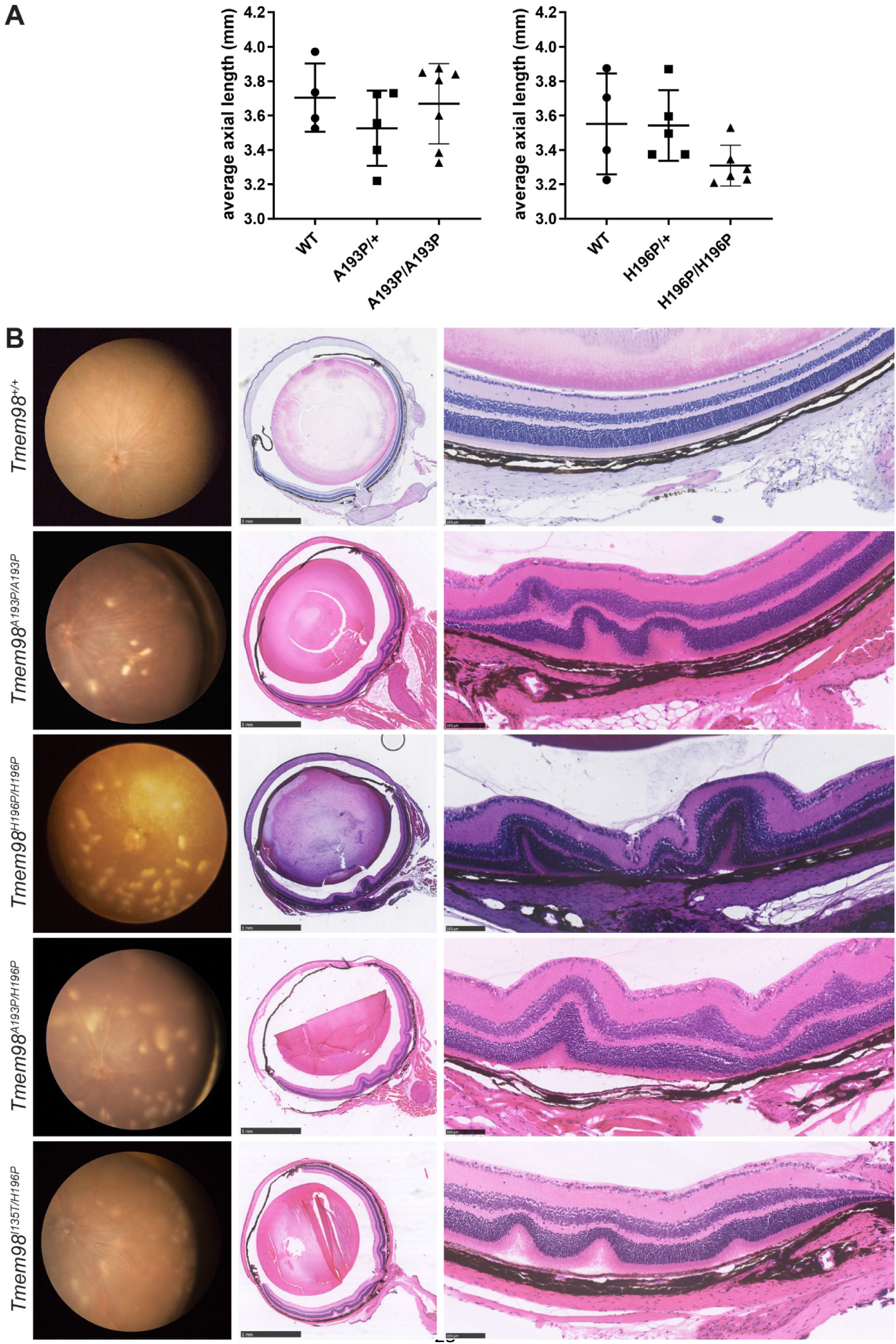
Eye phenotypes of homozygous and compound heterozygous mice with missense mutations of *Tmem98*. (**A**) Axial length measurements. Shown are the average axial length measurements for each mouse. (**B**) Left panel, retinal images; centre panel sections through the optic nerve; right panel, higher magnification pictures of the retina. *Tmem98*^*A193P/A193P*^, *Tmem98*^*H196P/H196P*^,*Tmem98*^*A193P/H196P*^ and *Tmem98*^*I135T/H196P*^ retinas all have scattered white spots (left panel) and folds in the outer nuclear layer and sometimes the inner retinal layers as well (centre and right panels). Scale bars: 1 mm (**B**, centre panel), 100 µM (**B**, right panel).

**Figure 3.**
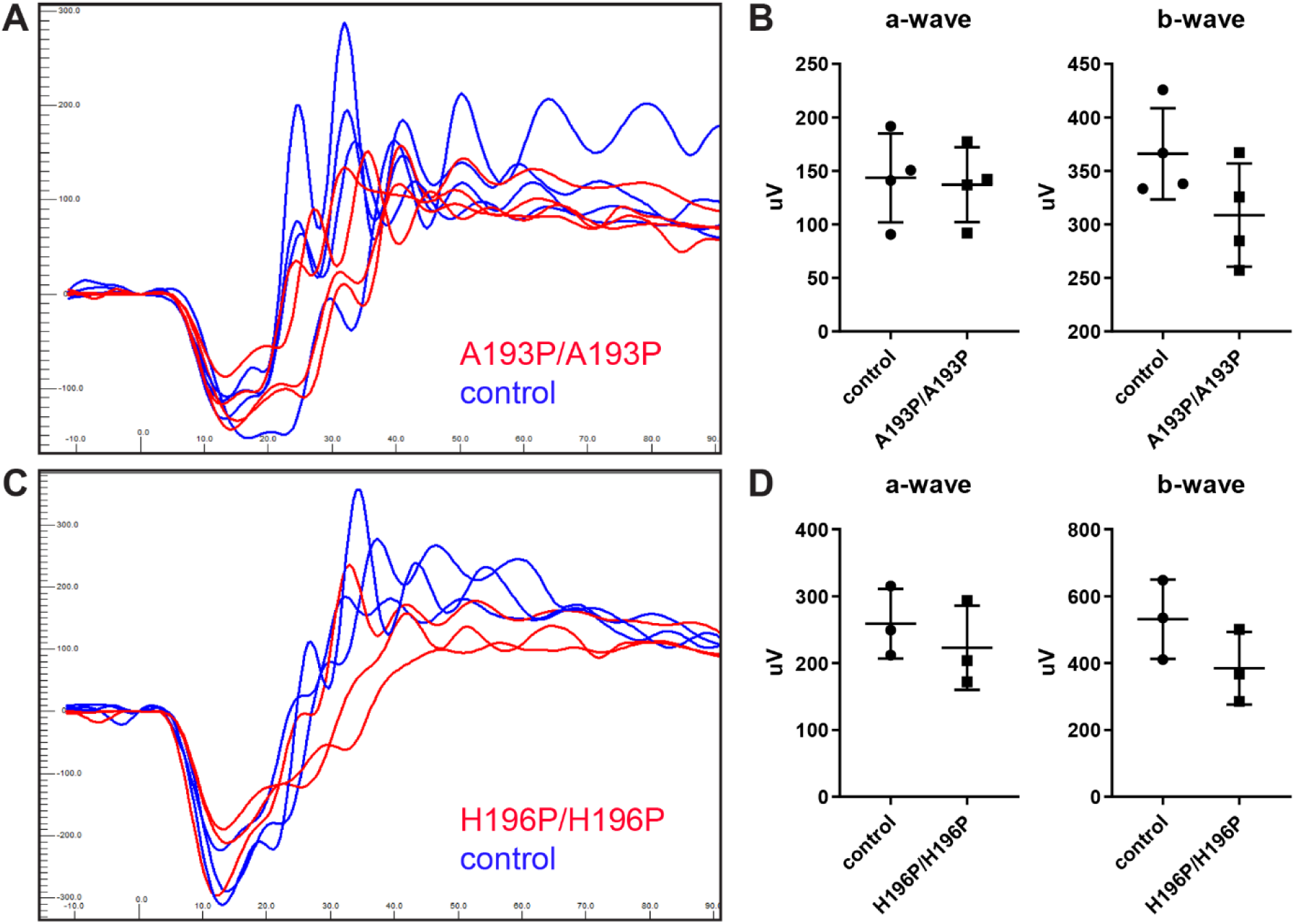
*Tmem98*^*A193P/A193P*^ and *Tmem98*^*H196P/H196P*^ mice have a normal ERG response. (**A-B**) Littermates, four *Tmem98*^*A193P/A193P*^ mice and four control mice (two wild-type and two *Tmem98*^*A193P/+*^) were tested at four months of age. (**A**) ERG traces of *Tmem98*^*A193P/A193P*^ mice (red lines), and control mice (blue lines). Shown are the responses at 3 cd.s/m^2^ (average of 4 flashes) for the left eye. (**B**) Comparison of a-wave amplitudes (right) and b-wave amplitudes (left), average of left and right eye for each mouse. There is no significant difference between *Tmem98*^*A193P/A193P*^ and control mice (a-wave, unpaired t test with Welch’s correction, P = 0.82 and b-wave, unpaired t test with Welch’s correction, P = 0.13). (**C**-**D**) Three *Tmem98*^*H196P/H196P*^ mice and three control mice (two wild-type and one *Tmem98*^*H196P/+*^) were tested at 6 months of age. (**C**) ERG traces of *Tmem98*^*H196P/H196P*^ mice (red lines), and control mice (blue lines). Shown are the responses at 3 cd.s/m^2^ (average of 4 flashes) for the left eye. (**B**) Comparison of a-wave amplitudes (left) and b-wave amplitudes (right), average of left and right eye for each mouse. There is no significant difference between *Tmem98*^*H196P/H196P*^ and control mice (a-wave unpaired t test with Welch’s correction, P = 0.49 and b-waveunpaired t test with Welch’s correction, P = 0.19).

## RESULTS

### *Rwhs* is a missense mutation of the *Tmem98* gene

The *N*-ethyl-*N*-nitrosourea (ENU)-induced mouse mutation retinal white spots (*Rwhs*) was found in a screen for dominant eye mutations ^37^. Mice heterozygous for the mutation have white patches on retinal imaging, apparently corresponding to folds or invaginations of the photoreceptor layers (Fig. 1A-B). Initial mapping indicated that *Rwhs* was located within an

8.5 Mb region of chromosome 11 ^37^. The retinal phenotype was found on a mixed Balb/c and C3H background. When *Rwhs* mutant mice were backcrossed to the C57BL/6J strain to refine the genetic mapping, some obligate heterozygous mice had retinas with a normal appearance, indicating that the dominant *Rwhs* phenotype is not completely penetrant and that modifiers in the C57BL/6J strain can attenuate it (Supplementary Fig. S1A-D). Crossing *Rwhs* to the wild-derived *CAST* strain to introduce a different and diverse genetic background, restored the retinal phenotype (Supplementary Fig. S1E-F). Intercrossing of heterozygous mice produced no homozygous offspring at weaning, whereas at late gestation (E17.5-E18.5) fetuses of the three expected genotypes were present at Mendelian ratios indicating that homozygous *Rwhs* is perinatally lethal (Table 2) (an initial report suggesting that homozygous *Rwhs* mice were viable was incorrect and due to mapping errors ^37^). We mapped this lethal phenotype by intercrossing recombinant animals and refined the critical interval to a 200 kb region between the single nucleotide polymorphism markers rs216663786 and rs28213460 on chromosome 11. This region contains the *Tmem98* gene and parts of the *Myo1d* and *Spaca3* genes. We amplified and sequenced all the exons and flanking regions from this region from *Rwhs* mutant mice along with controls. In addition we captured and sequenced all genomic DNA in the candidate interval. We found only a single nucleotide change in the mutant strain compared to the strain of origin, a T to C transition, in exon 5 of *Tmem98* (position 11:80,817,609 Mouse Dec. 2011 (GRCm38/mm10) Assembly (https://genome.ucsc.edu/)) (Fig. 1C). This mutation leads to the substitution of the non-polar aliphatic amino acid, isoleucine, by the polar amino acid threonine (I135T, numbering from entry Q91X86, http://www.uniprot.org). This amino acid substitution is predicted to be possibly damaging by the polymorphism prediction tool PolyPhen-2^42^.

TMEM98 is a 226 amino acid protein annotated with a transmembrane domain spanning amino acids 4-24 close to the N-terminus (http://www.uniprot.org). It is highly conserved across species; mouse and human TMEM98 share 98.7% amino acid identity and between mouse and *Ciona intestinalis*, the closest invertebrate species to the vertebrates, there is 38.6% amino acid identity in which I135 is conserved (Supplementary Fig. S2). TMEM98 has been reported to be a single-pass type II transmembrane protein in which the C-terminal part is extracellular ^49^. *TMEM98* is widely expressed and is reported to be most highly expressed in human and mouse RPE (http://www.biogps.org). We confirmed its high expression in the RPE and, within the retina, we also find expression at a lower level in the ganglion cell layer (Fig. 1D). To assess retinal function electroretinography was carried out on *Tmem98*^*I135T/+*^ and wild-type mice (Fig. 1E and F). There were no significant differences in the a-wave or b-wave amplitudes between the *Tmem98*^*I135T/+*^ and wild-type mice.

We also investigated the effect of loss-of-function of *Tmem98*. Heterozygous loss-of-function mice are viable and fertile and have normal retinas (Supplementary Fig. S6D and F).

Matings of heterozygous mice carrying the “knock-out first” *Tmem98*^*tm1a*^ allele produced no homozygous offspring (Supplementary Table S3) demonstrating that loss-of-function of *Tmem98* is lethal. At E16.5-E17.5 the three expected genotypes were present at Mendelian ratios (Supplementary Table S3) and in one litter collected at birth there were three homozgyotes and three wild-types indicating that lethality occurs between birth and weaning.

### The Human Nanophthalmos Missense Mutations Cause a Retinal Phenotype in the Mouse

Three mutations in *TMEM98* have been implicated in autosomal dominant nanophthalmos in human families ^12, 13^. Two are missense mutations, A193P and H196P. Both missense mutations affect amino acids that are highly conserved (Supplementary Fig. S2) Both are predicted to be probably damaging by the polymorphism prediction tool PolyPhen-2^42^.

To investigate the effect of the two missense mutations we used CRISPR-Cas9 to introduce A193P and H196P into the mouse gene and established lines carrying each. Western blot analysis using a validated anti-TMEM98 antibody (Supplementary Fig. S3A) showed that the mutant proteins are expressed (Supplementary Fig. S3B). Heterozygous mice for both missense mutations were viable and fertile and did not exhibit any gross eye or retinal defects when examined between 5-9 months of age (Supplementary Fig. S4, *Tmem98*^*A193P/+*^, n=10; *Tmem98*^*H196P/+*^, n=19). In contrast to the *Tmem98*^*I135T*^ and knock-out alleles, homozygotes for both the *Tmem98*^*A193P*^ and *Tmem98*^*H196P*^ alleles were viable and found at the expected Mendelian ratios (Supplementary Table S4). The eyes of homozygous mice do not appear to be significantly different in axial length when compared to wild-type eyes (Fig. 2A). Although the mean length of homozygous *Tmem98*^*H196P*^ eyes is ~7% smaller than controls this does not have statistical support (P=0.1). From about 3 months of age we found that white patches developed on the retinas of the homozygous mice and on histological examination we found folds or invaginations in the retinal layers (Fig. 2B). The appearance of the white patches was progressive; at younger ages the retinas of some homozygous mice appeared normal with patches becoming apparent as they aged (Supplementary Fig. S5). In the A193P line only 6 of 7 homozygotes were found to have retinal defects at 6 months of age; the seventh developed white patches on the retina by 9 months. In the H196P line 4/20 homozygous mice that were examined between 3 and 3.5 months of age appeared to have normal retinas. We crossed the lines together to generate compound heterozygotes. All *Tmem98*^*A193P/H196P*^ (n=4), *Tmem98*^*I135T/A193P*^ (n=7) and *Tmem98*^*I135T/H196P*^ (n=5) mice examined also displayed a similar phenotype of white patches on the retina (Fig. 2B and data not shown). We also crossed *Tmem98*^*H196P*^ mice with mice carrying a loss-of-function allele *Tmem98*^*tm1b*^. Compound heterozygous mice were viable and of 17 mice examined all had normal retinas except for one mouse which had three faint spots on one retina at one year of age (Supplementary Fig. S6). These results suggest that a threshold level of the mutant missense TMEM98^H196P^ and TMEM98^A193P^ proteins is required to elicit the formation of white patches on the retina and that the missense mutations found in the human nanophthalmos patients are not loss-of-function. Furthermore, mice heterozygous for *Tmem98*^*tm1b*^ with a wild type allele do not have the retinal phenotype (Supplementary Fig. S6).

To assess retinal function electroretinography was carried out on *Tmem98*^*A193P/A193P*^,*Tmem98*^*H196P/H196P*^ and control mice (Fig. 3). There were no significant differences in the a-wave or b-wave amplitudes between the control and the mutant mice. To determine if there was any deterioration in retinal function in older mice we also tested *Tmem98*^*H196P/H196P*^and control mice at 9-11 months of age but again we did not find any significant differences in the a-wave or b-wave amplitutes between the control and the mutant mice (Supplementary Fig. S7).

### Characterisation of the Retinal Folds Caused by the A193P and H196P Mutations

We investigated the retinal folds by immunostaining of retinal sections. We found that in the interior of retinal folds the outer segment layer is massively expanded as demonstrated by positive staining for the transmembrane protein prominin-1 and the rod and cone opsins (rhodopsin and ML opsin), indicating that the folds are filled with excess, disorganised outer segments or remnants of outer segments (Fig. 4).

**Figure 4.**
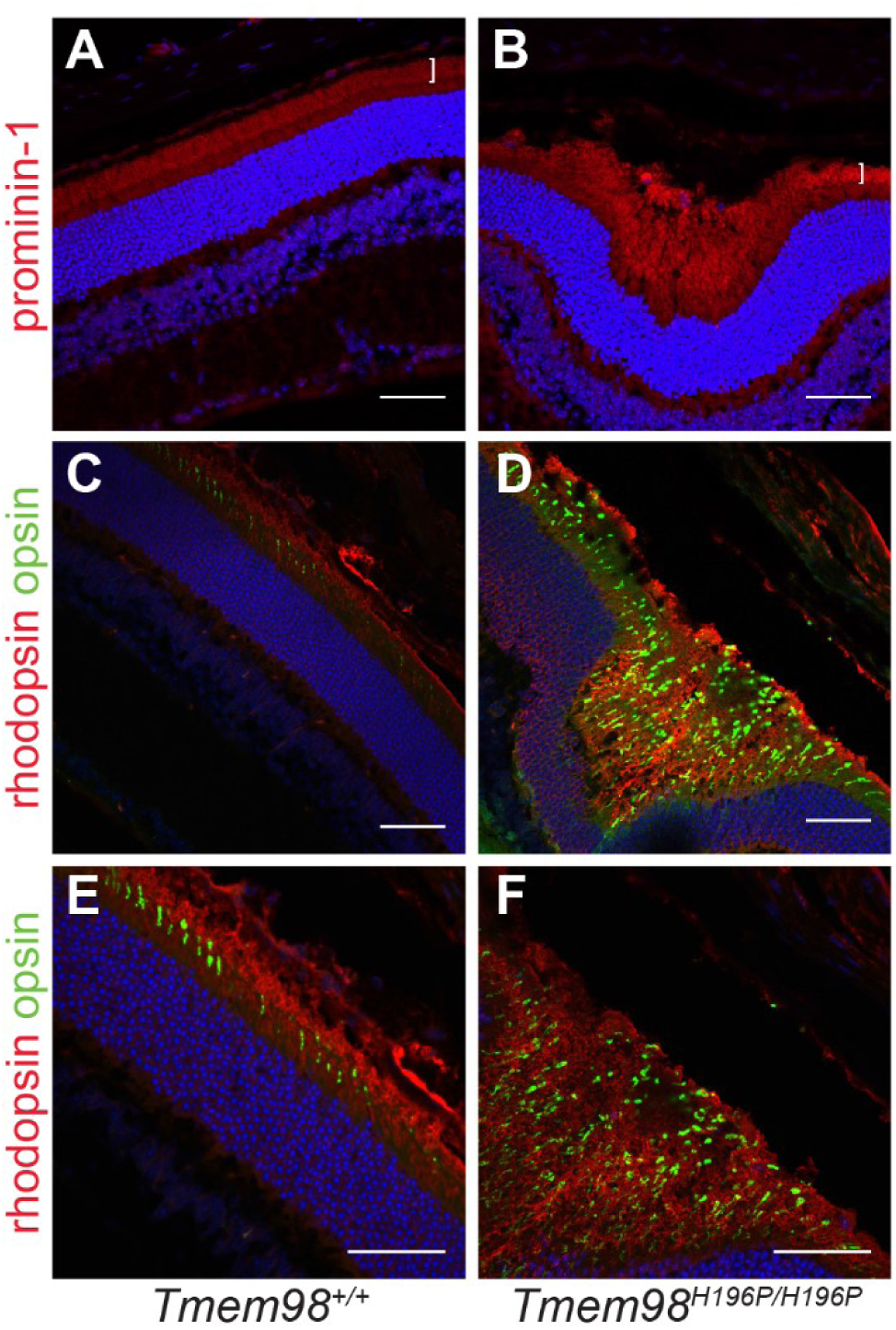
The interiors of the retinal folds found in the *Tmem98*^*H196P/H196P*^ mutant mice are filled with excess outer segments. Immunostaining of retinal sections from wild-type mice (**A**, **C** and **E**) and *Tmem98*^*H196P/H196P*^ mice (**B**, **D** and **F**). (**A**-**B**) Prominin-1 staining (red) shows that the outer segment layer (white bracket) is expanded in the retinal fold of the mutant (**B**) compared to wild-type (**A**). (**C**-**F**) Rhodopsin (red) and opsin (green) staining shows that the interior of the retinal fold is filled with outer segments. DAPI staining is shown in blue. Scale bars: 50 µM.

The retinal folds seen in other mutant mouse lines are accompanied by defects in the OLM. This structure is formed by adherens junctions between the Müller glia cells and the photoreceptors. We investigated the integrity of the OLM in our mutant mice by staining for β-catenin (Fig. 5A-D). In control mice and in the regions of the retinas of mutant mice unaffected by folds the OLM appeared intact. However, at the folds the OLM is clearly disrupted and gaps can be seen suggesting that cell-cell connections have been broken. Reactive gliosis indicated by upregulation of GFAP in the Müller cells is a response to retinal stress, injury or damage ^50^. We observed abnormal GFAP staining in the mutant retinas that was confined to the regions with folds, indicating that in these areas, but not elsewhere in the retina, there is a stress response (Fig. 5E-H, Supplementary Fig. S8A-D). We also stained for F4/80, a marker for macrophages/microglia. In the mutant retinas positive staining was found in the interior of retinal folds but not elsewhere in the photoreceptor layer (Fig. 5J-L, Supplementary Fig. S8F-H). The amoeboid shape of the positively-stained cells suggests that they are macrophages that have infiltrated into the retinal folds containing excess and disorganised outer segments and that they are phagocytosing degenerating outer segments. We did not observe melanin within the macrophages indicating that they had not engulfed pigmented RPE cells. Finally, we examined by transmission electron microscopy the ultrastructural architecture of the boundary between the RPE and outer segments (Fig. 6). For the homozygous mutants, in retinal areas without folds, the boundary between outer segments and RPE appeared normal (compare Fig. 6A with Fig. 6B and D). However, in the areas with folds the outer segments were disorganised and appeared to be degenerating, with cavitites and other celluar debris apparent (Fig. 6C and 6E).

**Figure 5.**
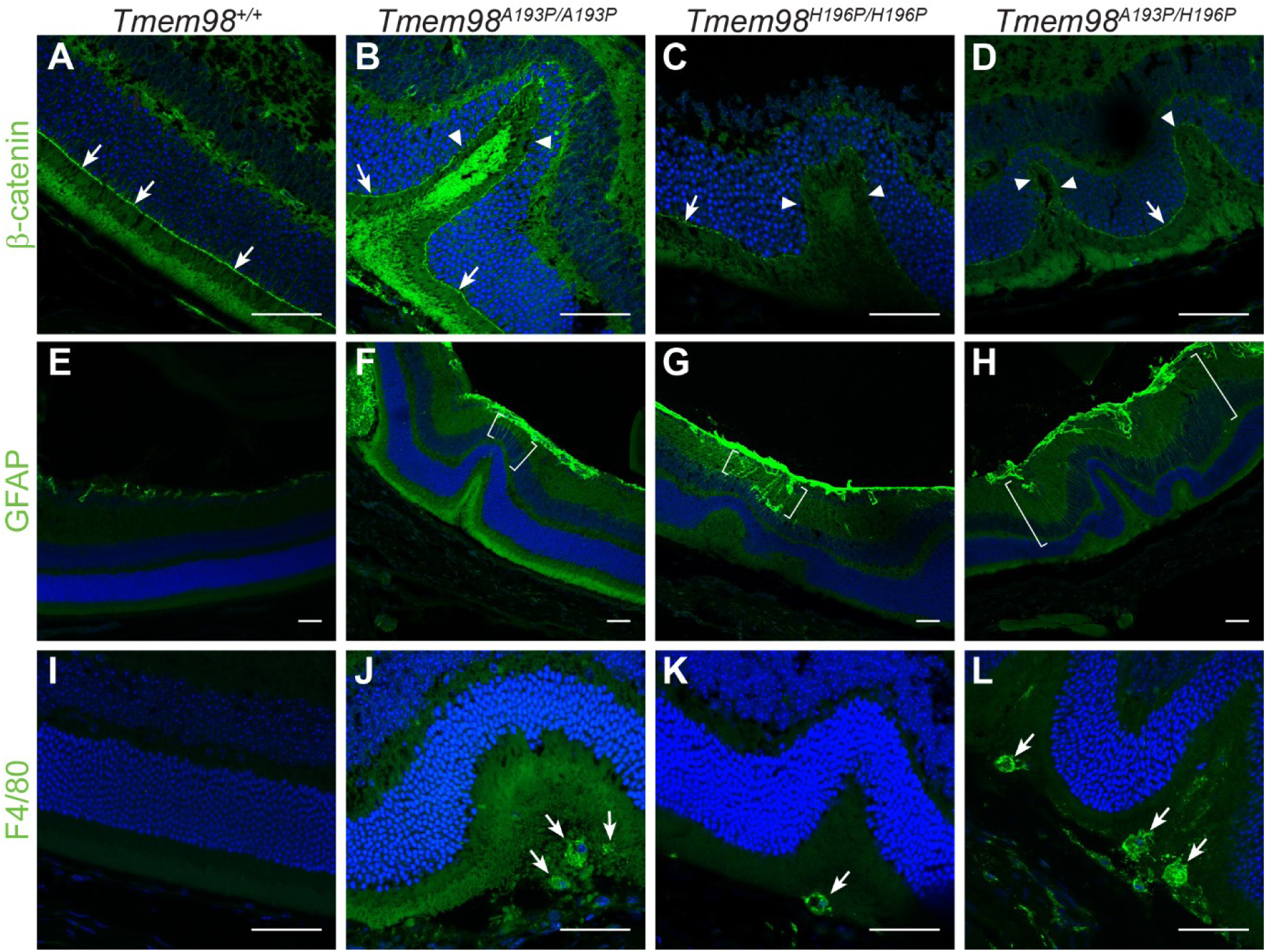
Characterisation of the retinal folds in the *Tmem98* mutant mice. Immunostaining of retinal sections from wild-type mice (**A**, **E** and **I**), *Tmem98*^*A193P/A193P*^ mice (**B**, **F** and **J**), *Tmem98*^*H196P/H196P*^ mice (**C**, **G** and **K**), *Tmem98*^*A193P/H196P*^ mice (**D**, **H** and **L**). (**A**-**D**) β-catenin staining (green) shows that the OLM is intact in the wild-type retina and in the areas either side of folds in the mutant retinas (white arrows) but in the folds of the mutant retinas the OLM is interrupted (white arrowheads). (**E**-**H**) GFAP staining (green) is normal in the wild-type retina but above the folds in the mutant retinas there is abnormal GFAP staining extending towards the outer nuclear layer (areas between the white brackets). This indicates that in the mutants the retina is stressed in the folded regions and that retinal stress is confined to the folds. (**I**-**L**) F4/80 staining (green) reveals that macrophages have infiltrated into the areas below the folded outer nuclear layer containing excess photoreceptors in the mutant retinas (white arrows). Staining was not observed outside the folds in the mutant retinas. DAPI staining is shown in blue. Scale bars: 50 µm.

**Figure 6.**
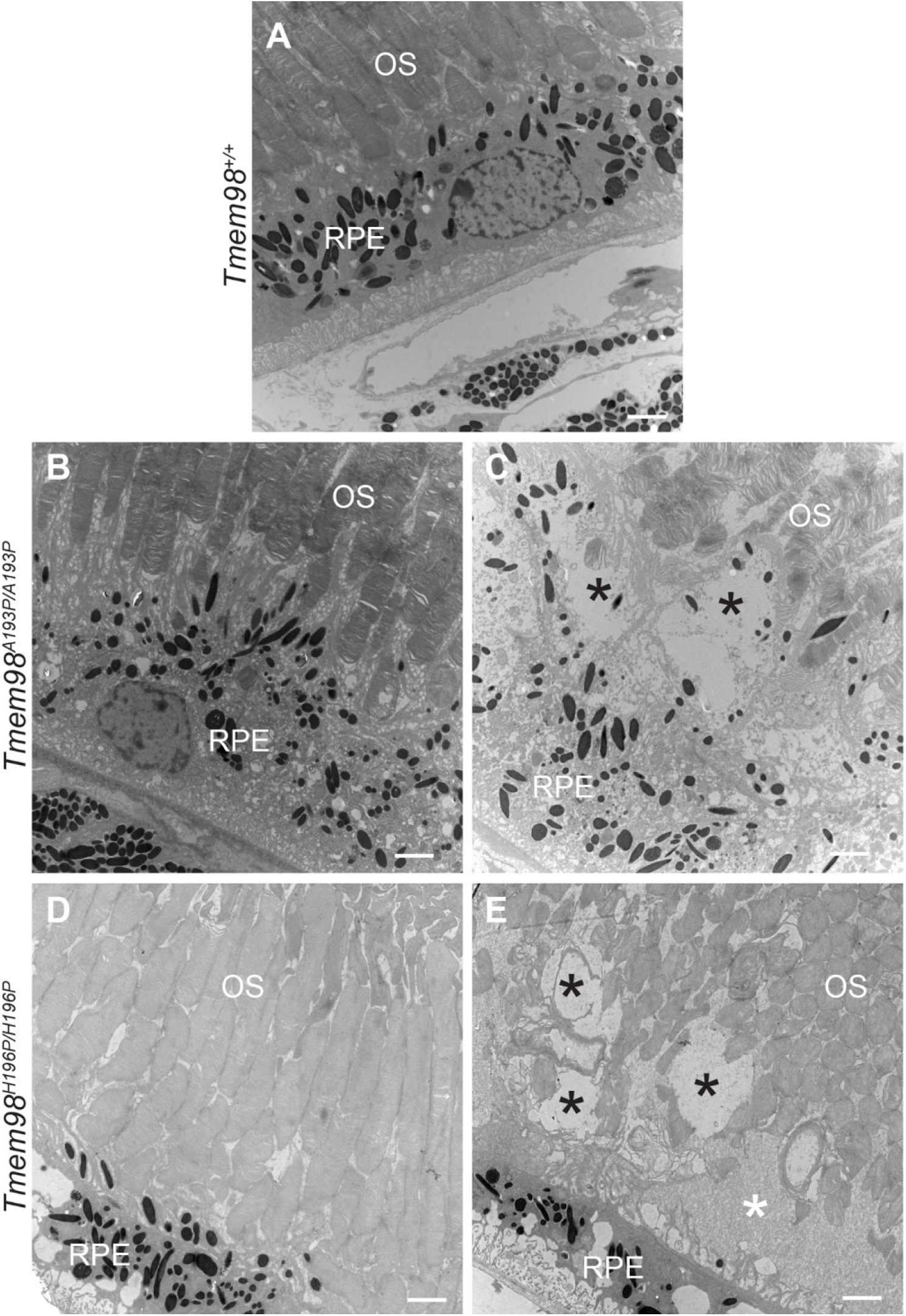
Ultrastructural analysis of the RPE and outer segment boundary. (**A**) Wild-type mice are normal. (**B** and **D**) In mutant mice from retinal regions with no folds the outer segments adjacent to the RPE appear normal. (**C** and **E**) In retinal areas with folds the outer segments abutting the RPE appear abnormal and disorganised. Several large vacuoles (indicated by black asterisks) can be seen. In (**E**) there is an area containing cellular debris (indicated by a white asterisk). *Tmem98*^*A193P/A193P*^ (**B**, **C**) and *Tmem98*^*H196P/H196P*^ (**D**, **E**). OS = outer segments, RPE = retinal pigment epithelium. Scale bars: 2µm.

## DISCUSSION

### *Rwhs* is Caused by a Missense Mutation in *Tmem98* that is Homozygous Lethal

Here we report that the ENU-induced dominant retinal white spotting phenotype, *Rwhs*, is caused by an I135T missense mutation in the highly conserved transmembrane protein encoding gene *Tmem98*. We also found that when homozygous the *Tmem98*^*I135T*^ allele is perinatally lethal. *Tmem98* was one of the genes included in an international project to produce and phenotype knockout mouse lines for 20,000 genes ^51^. The targeted allele, *Tmem98*^*tm1a*^, was subjected to a high-content phenotyping pipeline (results available at http://www.mousephenotype.org/data/genes/MGI:1923457). It was found to be lethal pre-weaning as homozygotes, but no significant heterozygous phenotypic variation from wild-type was reported. Neither their slit lamp analysis nor our retinal examination found any eye defects in knock-out heterozygous mice (Supplementary Fig. S6D and F). We also found that *Tmem98*^*tm1a*^ is homozygous lethal and narrowed the stage of lethality to the perinatal stage (Supplementary Table S2). This suggests that haploinsufficency for TMEM98 protein does not cause a retinal phenotype and that the I135T mutation in TMEM98 is not a loss-of-function allele but changes the protein’s function leading to the retinal white spotting phenotype.

### Phenotype of Human Nanophthalmos Associated *TMEM98* Missense Mutations in the Mouse

*Tmem98* has been previously suggested to be a novel chemoresistance-conferring gene in hepatoceullular carcinoma ^52^. It has also been reported to be able to promote the differentiation of T helper 1 cells and to be involved in the invasion and migration of lung cancer cells ^49, 53^. Recently it has been reported that TMEM98 interacts with MYRF and prevents its autocatalytic cleavage ^54^. In relation to human disease two TMEM98 missense mutations, A193P and H196P, have been reported to be associated with dominant nanophthalmos ^12, 13^. We introduced these mutations into the mouse gene and found that mice homozygous for either, or compound heterozygous for both, developed white patches on their retinas accompanied by retinal folds, replicating the dominant phenotype found in the ENU-induced allele. We also observed for all three alleles separation of the RPE away from the neural retina in the areas under the folds. Mice lacking *Tmem98* do not survive after birth (Supplementary Table S3). The three missense mutations described here are not null. The two human-equivalent mutations in the mouse are homozygous viable (Supplementary Table S4), and the mouse mutation, when in combination with a knock-out allele is also viable (data not shown). Furthermore, the H196P missense mutation in combination with a gene knockout has no detectable eye phenotype and the mice are viable (Supplementary Fig. S6). This indicates that the recessive mutations are not loss of function, but are gain of function with, in the case of H196P at least, a threshold dosage requirement. However, the phenotypes are different from the reported dominant nanophthalmos caused by the same mutations in humans.

Other genes causing nanophthalmos may also be gain of function. The serine protease *PRSS56* is a recessive posterior microphthamia or nanophthalmos mutant gene in humans, with both identified mutations affecting the C-terminus of the protein ^8^. The mouse mutant model of this gene, which produces a variable and slight reduction in ocular length, is a splice site mutation resulting in a truncated protein which nevertheless has normal protease activity *in vitro* ^8^. Recently the phenotype of a null allele of *Prss56* has been described and it does cause some reduction in ocular size and hyperopia ^36^.

The mouse model of the nanophthalmos gene, *Mfrp*, does not reproduce the human phenotype. Rather, loss of function of this gene results in white spots apparent on retinal imaging (retinitis punctata albicans) but these have different origin from the phenotype we observe, and progress to photoreceptor degeneration. It has been reported that the *Mfrp* knockout mice have eyes that are slightly (but not statistically significantly) smaller, by about 2% in axial length^35^. Our analysis of the human-equivalent mouse *Tmem98* mutations show that that neither have statistically significant size reductions, although the H196P line is ~7% shorter It is worth noting that different strains of mice have measurably different ocular size. Strain differences of up to 2% and sex differences of over 1.5% have been reported^48^.

Two genes that are infrequently associated with nanopthalmos, *CRB1* and *BEST1*, both can produce a mouse phenotype apparently indistinguishable from the one we describe here. Furthermore, the knockout mouse model of the nuclear receptor gene, *Nrl*, also develops retinal folds during post-natal life^55^. The *Nrl* mutant eye has defects in the outer limiting membrane (OLM), a component of which is the CRB1 protein. The *Tmem98* mutations have OLM defects, but we cannot ascertain whether these defects are the primary cause of the folds or a secondary consequence.

Retinal folds are a characteristic of nanophthalmic eyes and this is usually attributed to a differential growth of neural retina within the smaller optic cup. Our data show that folds can be seen in a normal-sized eye, but we do not know whether there is nevertheless excess growth of the neural retina. The retinal defects we see are not associated with an ERG deficit, suggesting that the rest of the retina, unaffected by the folds, is functionally normal.

Notable is the detachment of the retina from the RPE within the folds. In the *Nrl* and *Crb1* mutant eyes the photoreceptors that have lost their connection to the RPE can be seen to degenerate. We see no evidence of photoreceptor degeneration in the *Tmem98* mutants. The mechanism of the pathology is still unclear. As *Tmem98* is strongly expressed in the RPE, and not at all in the photoreceptors, it is likely that RPE is the affected tissue. The key observation in these and other mouse models of nanophthalmos is that the defects are focal and progressive, suggesting that secondary events exacerbate an underlying but (given ERG data) non-pathological defect. The accumulation of photoreceptor cell debris below the retinal folds suggests a focal defect in outer segment phagocytosis but one that does not lead to photoreceptor degeneration. Three of the genes associated with nanophthalmos are expressed in the RPE; *Tmem98*, *Best1* and *Mfrp*. *Crb1* is expressed in photoreceptors and *Prss56*, a secreted serine protease, is expressed in the Müller cells. It is possible that these genes interact and affect a common pathway, indeed upregulation of *Prss56* has been observed in *Mfrp* mutants ^56^.

### Effect on TMEM98 Protein Structure of the Missense Mutations

TMEM98 has structural homology to Cyclin-D1-binding protein 1 (CCNDBP1) and contains a Grap2 and cyclin-D-interacting (GCIP) domain (pfam13324) spanning amino acids 49-152. Based upon the crystal structure of CCNDBP1 following the N-terminal transmembrane domain TMEM98 is predicted to have a five helix bundle structure using PHYRE2^44^ (Fig. 7A). I135 is located in the third α-helix completely buried within a sterically close-packed hydrophobic core of the domain where it has hydrophobic interactions with amino acids from four of the α-helices (Fig. 7B). SCWRL^46^ and FoldX^47^ stability calculations indicate that the *Rwhs* mutation T135 is only very mildly destabilising in this homology model. Its side-chain does not show steric clashes with other neighbouring atoms and its buried hydroxyl group forms a favourable hydrogen bond with the main-chain oxygen atom of A132. The stability energy calculation on the mutant TMEM98 domain structure (mean ΔΔG) is 0.28 kcal/mol (>1.6 kcal/mol is considered destabilising). The genetic evidence shows that the I135T is not loss-of-function. The detrimential effect of a mutation leading to loss of thermodynamic stability can be compensated for by the creation of novel functions^57^. This might be the case with the I135T mutation. The two missense mutations that are associated with nanophthalmos introduce a proline in the middle of the final α-helix of the protein which would likely lead to disruption of the secondary structure of the protein^58^. The recent finding that TMEM98 binds to and prevents the self-cleavage of the oligodendrocyte transcription factor MYRF^54^ (also highly expressed in the RPE; http://www.biogps.org) will perhaps lead to a mechanistic explanation, although the part of TMEM98 that binds to MYRF was mapped to the N-terminal 88 amino acids and a region between 55-152 amino acids was shown to be required for the cleavage inhibition. As the nanophthalmos patient-specific missense mutations are more C-terminal their effect on this reported interaction will be the subject of future work.

**Figure 7.**
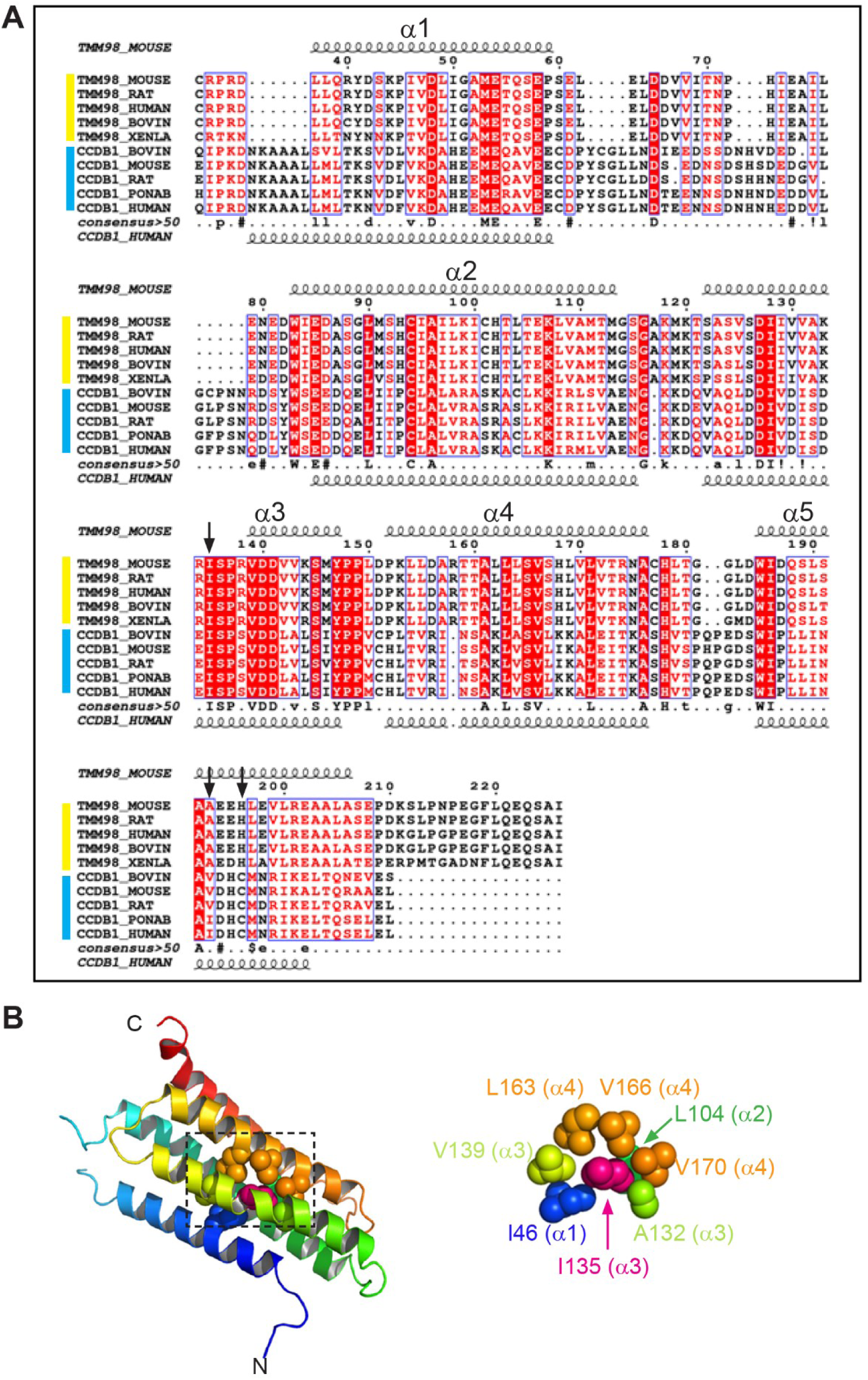
TMEM98 protein structure and predicted effect of the I135T mutation (**A**) Alignment of TMEM98 proteins (yellow group) and CCNDBP1 proteins (blue group). The positions of predicted α-helices (1-5) are indicated by the curly lines. The positions of the three missense mutations are indicated by the black arrows. (**B**) Helical bundle based upon the crystal structure of CCNDBP1 (c3ay5A). The N- and C-terminal ends are indicated. The region surrounding I135 (dashed box) is enlarged on the right showing the hydrophobic interactions within 5 Å of I135.

## Acknowledgements

We thank Morag Robertson for help with genotyping, Craig Nicol and Connor Warnock for help with photography, the IGMM Advanced Imaging Resource for help with imaging, Edinburgh University Bioresearch and Veterinary Services for animal husbandry and MRC Human Genetics Unit scientific support services.

## Supplementary Methods

### Mice

By crossing *Tmem98*^*tm1a*/+^ mice with mice carrying Flpe the *tm1a* ‘knockout-first’ allele was converted to a conditional allele *Tmem98*^*tm1c(EUCOMM)Wtsi*^ (hereafter *Tmem98*^*tm1c*^). By crossing *Tmem98*^*tm1c*/+^ mice with mice carrying Cre the floxed critical exon 4 is deleted generating a deletion allele that has a frame shift *Tmem98*^*tm1d(EUCOMM)Wtsi*^ (hereafter *Tmem98*^*tm1d*^) that would be subject to nonsense mediated decay ^39^.

### Embryonic Protein Preparations

Embryonic day 12.5 (E12.5) embryos were collected and a small piece of tail used for genotyping. Tissue was homogenised in RIPA buffer (Cell Signaling Technology) plus 1 mM phenymethysulfonyl (ThermoFisher Scientific) and Complete Protease Inhibitor Cocktail (Roche) and sonicated for 30s 3 times. Crude lysates were cleared by centrifugation (20,000 g for 30 minutes at 4°C) and protein concentrations determined by Bradford assay (Bio-Rad). 20 µg samples were used for Western blotting.

### Western Blotting

Equal amounts of protein lysates were separated on 4-12% Nupage Bis-Tris gels (ThermoFisher Scientific) and transferred to polyvinylidene difluoride or nitrocellulose membranes. Membranes were blocked for one hour at room temperature in SuperBlock T20 (TBS) Blocking Buffer (ThermoFisher Scientific) and incubated with primary antibodies for one hour at room temperature or overnight at 4°C in blocking buffer with shaking. Following washing with TBST membranes were incubated with ECL horse radish peroxidase (HRP)-conjugated secondary antibodies (GE Healthcare) diluted 1:5000 in blocking buffer for one hour at room temperature, washed with TBST and developed using SuperSignal™ West Pico PLUS (ThermoFisher Scientific).

## Supplementary Tables

**Supplementary Table S1.**
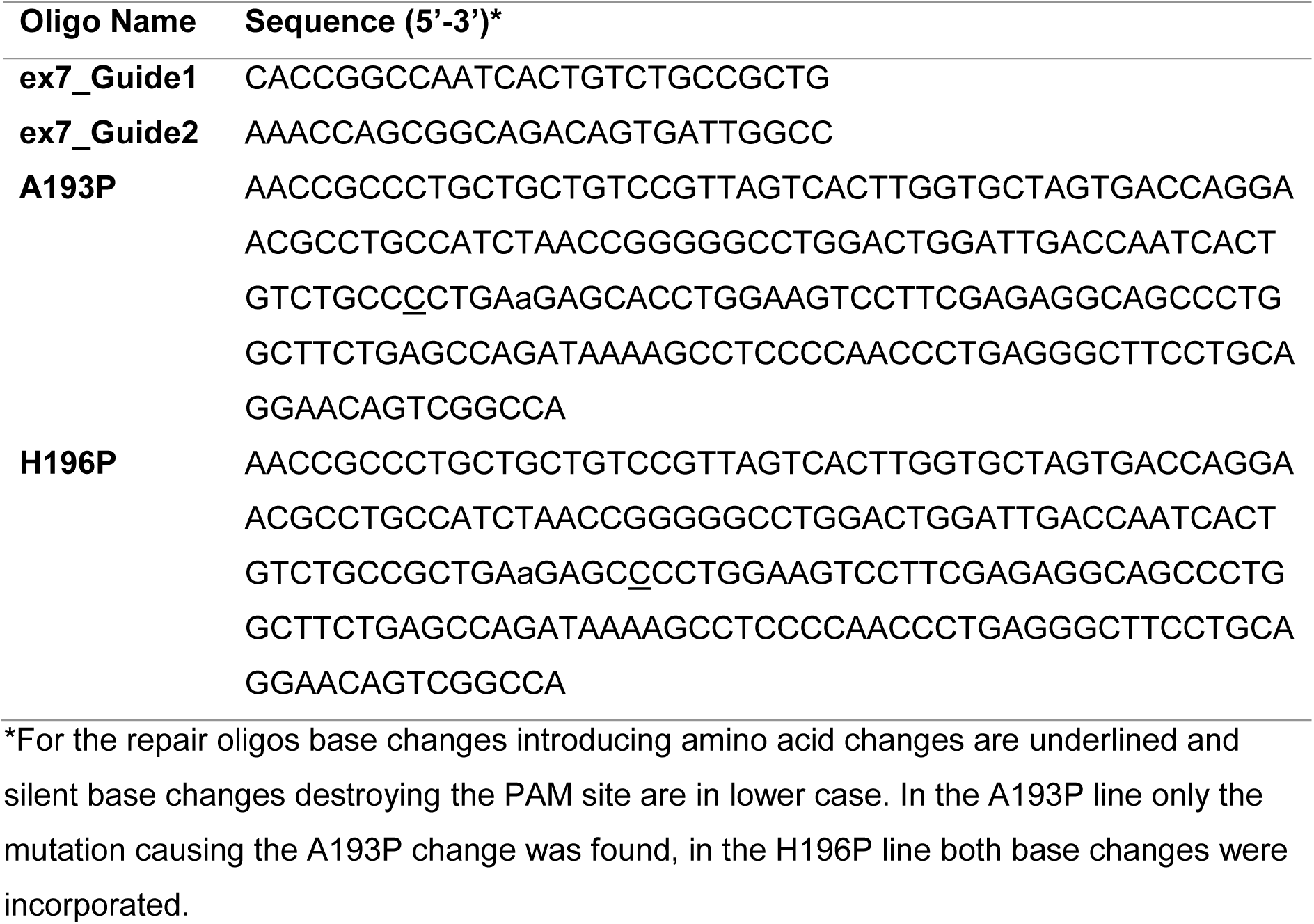
Sequences of oligos

**Supplementary Table S2.**
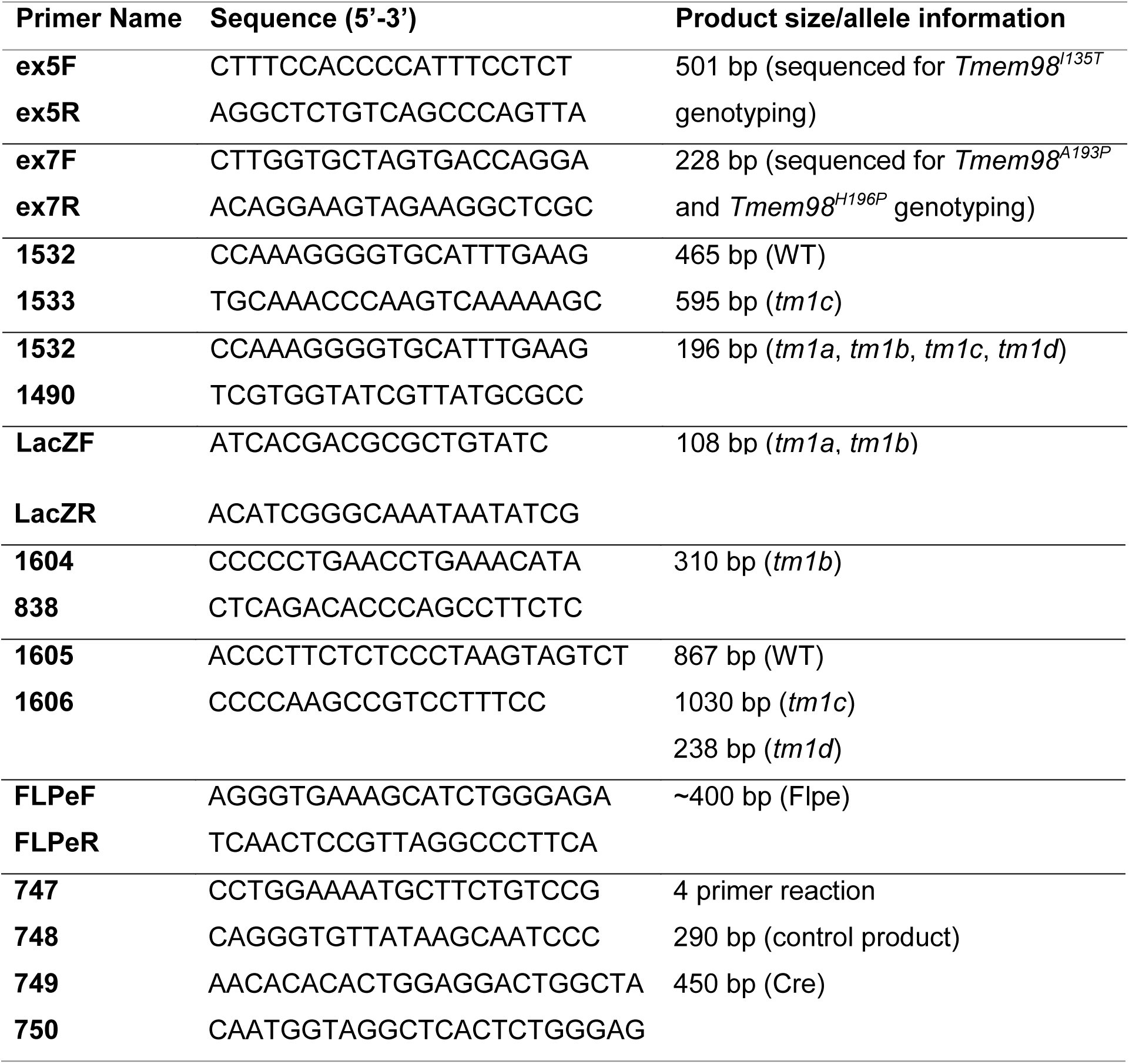
Genotyping primers

**Supplementary Table S3.** *Tmem98*^*tm1a/+*^ intercross genotyping results

| **Age** | **WT** | ***tm1a*/+** | ***tm1a*/*tm1a*** | **Total** | **P\*** |
| --- | --- | --- | --- | --- | --- |
| at weaning | 21 | 38 | 0 | 59 | <0.0001 |
| E16.5-E17.5 | 10 | 20 | 10 | 40 | 1.0000 |

\*Test for significance using chi-square test

**Supplementary Table S4.** Mouse homologues of the human missense nanophthalmos *TMEM98* mutations intercross genotyping results at weaning

| Cross | WT | A193P/+ | A193P/A193P | H196P/+ | H196P/H196P | A193P/H196P | Total | P* |
| --- | --- | --- | --- | --- | --- | --- | --- | --- |
| A193P/+ x A193P/+ | 10 | 21 | 12 | N/A | N/A | N/A | 43 | 0.9006 |
| H196P/+ x H196P/+ | 68 | N/A | N/A | 94 | 49 | N/A | 211 | 0.0516 |
| A193P/+ x H196P/+ | 23 | 18 | N/A | 20 | N/A | 14 | 75 | 0.5164 |

\*Test for significance using chi-square test N/A=not applicable

**Supplementary Figure S1.**
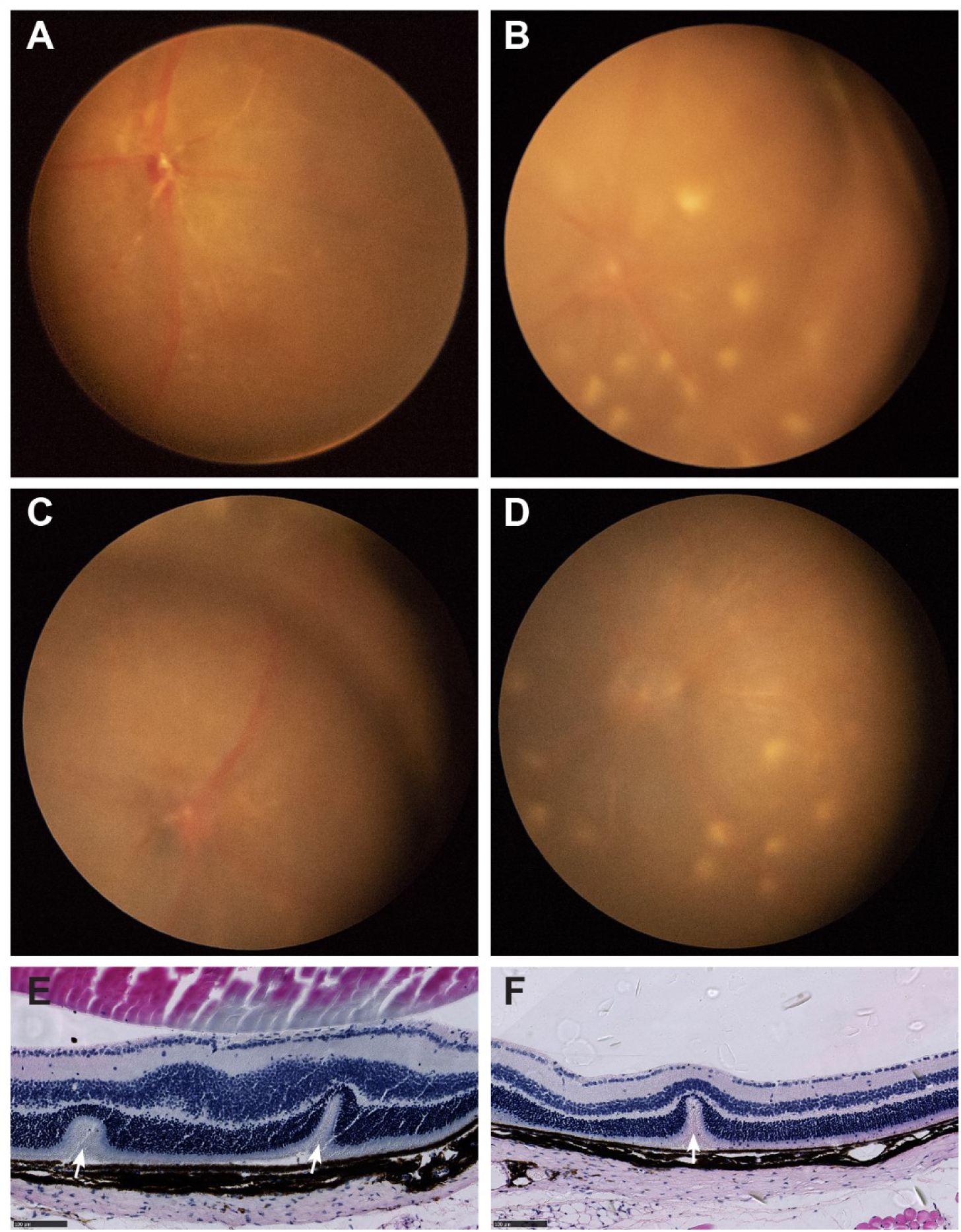
Effect of genetic background on the *Rwhs* dominant retinal phenotype. (**A**-**D**) Retinal images of *Tmem98*^*135T/+*^ mice on the C57BL/6J genetic background. **A** and **B** are littermates and **C** and **D** are littermates. The mice shown in **A** and **C** have normal retinas whereas the mice shown in **B** and **D** have white spots on their retinas. (**E**-**F**) Haematoxylin and eosin stained sections of *Tmem98*^*135T/+*^ retinas on the CAST strain background displaying invaginations of the outer nuclear layer indicated by arrows. Scale bars: 100 µm.

**Supplementary Figure S2.**
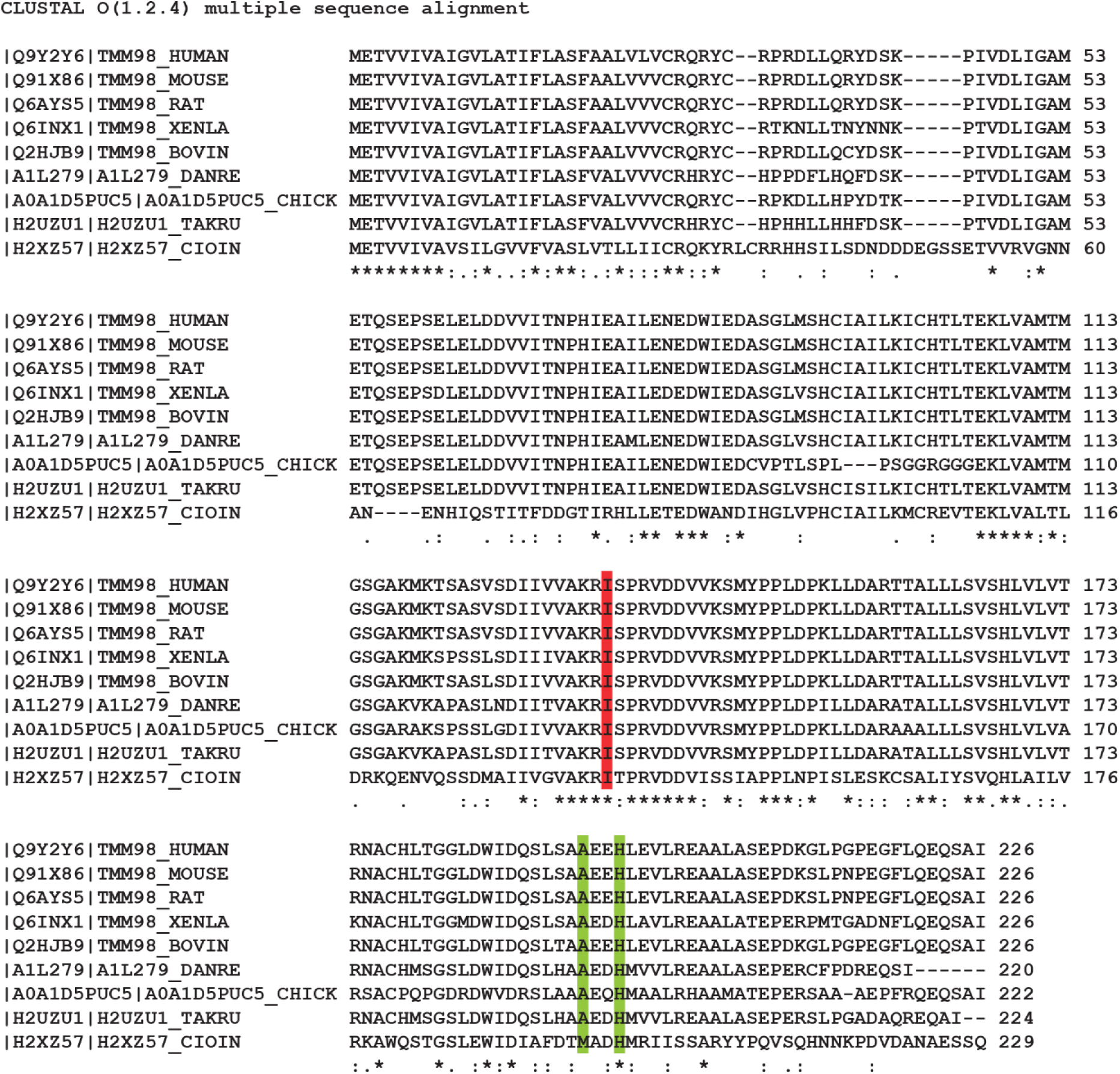
TMEM98 protein sequences from different species aligned using the Clustal Omega program. The uniprot accession numbers and identifiers are shown on the left. An * indicates a completely conserved residue, a : indicates that residues have strongly similar properties (scoring > 0.5 in the Gonnet PAM 250 matrix), a. indicates that residues have weakly similar properties (scoring =< 0.5 in the Gonnet PAM 250 matrix). I135, which is mutated in *Rwhs*, is completely conserved and highlighted in red. The two residues affected in the human nanophthalmos patients, A193 and H196, are highlighted in green. Both are completely conserved except that a methionine is substituted for A193 in *Ciona intestinalis*.

**Supplementary Figure S3.**
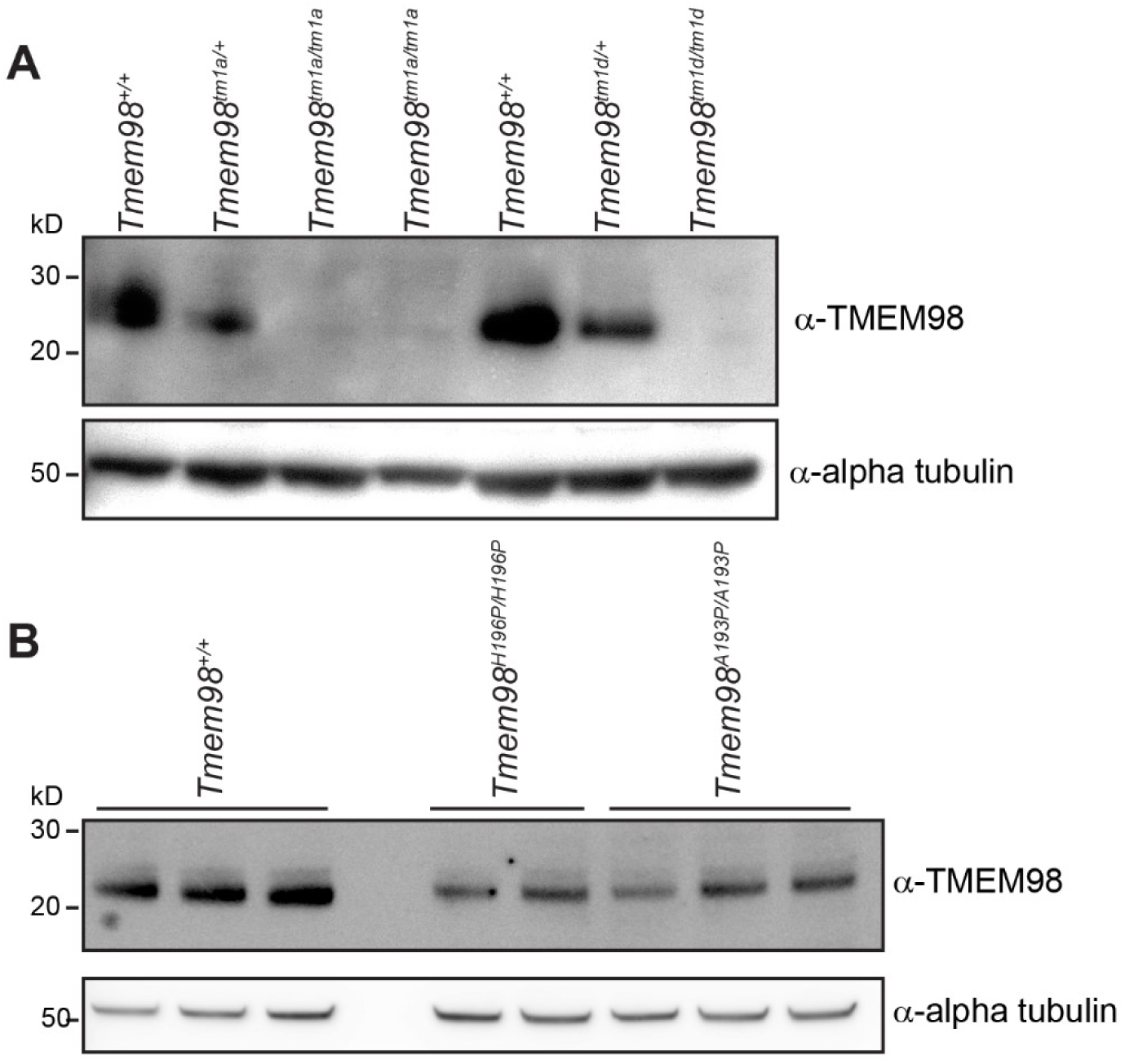
TMEM98 protein is expressed in the missense mutants. (**A**) Western blot analysis of E12.5 embryonic protein lysates of the indicated genotypes. TMEM98 protein can be detected in the wild-type and heterozygous samples but not the homozygous knock-out samples validating the antibody. (**B**) Western blot analysis of E12.5 embryonic protein lysates of the indicated genotypes. TMEM98 is present in the homozygous mutants carrying the missense mutations found in human nanophthalmos patients. Alpha tubulin was used as a loading control.

**Supplementary Figure S4.**
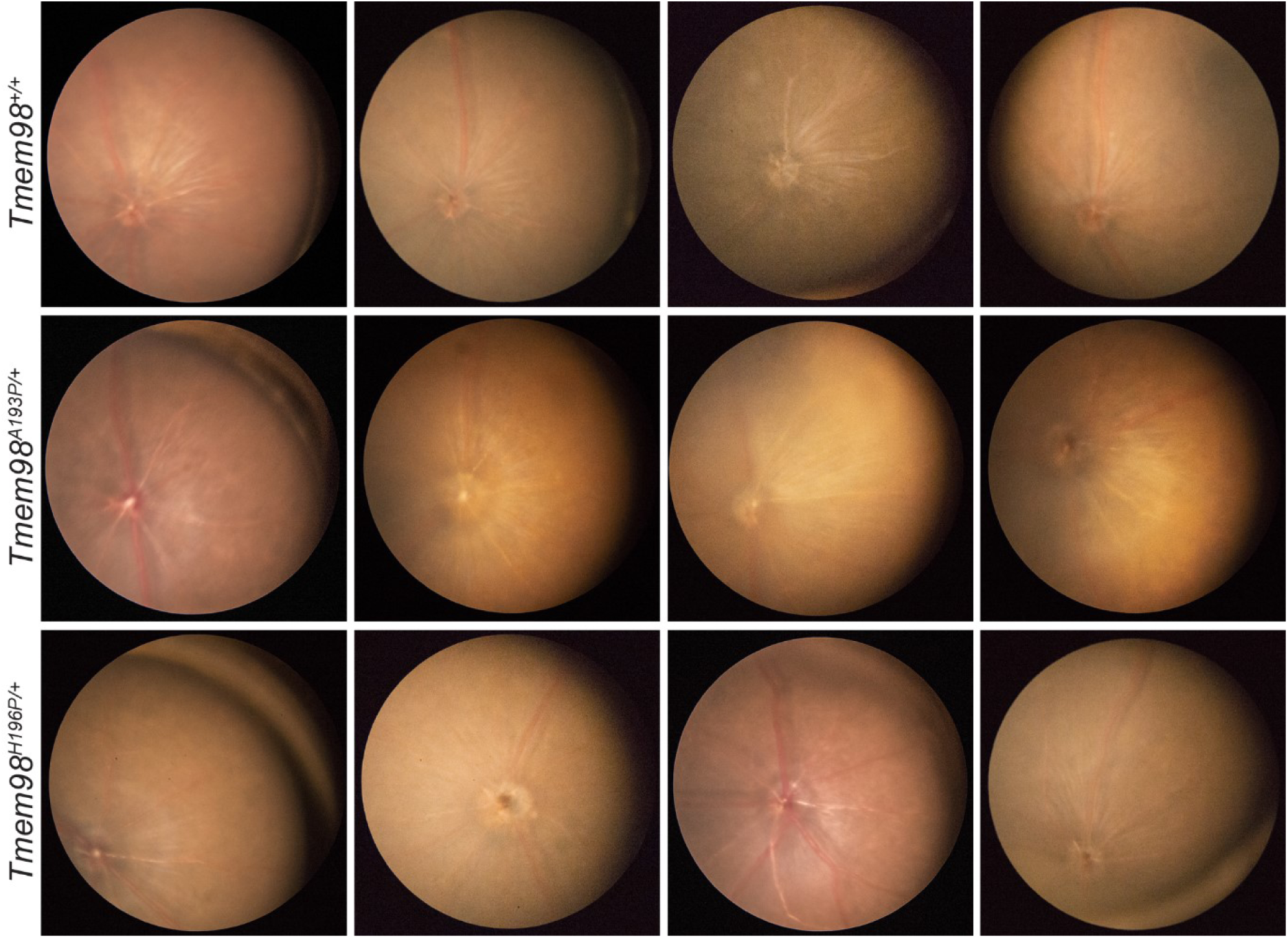
Retinas of mice heterozygous for the human nanophthalmos missense mutations are normal. Retinal pictures from mice of between 5-10 months are shown. Top row wild-type mice, middle row *Tmem98*^*A193P/+*^ mice and bottom row *Tmem98*^*H196P/+*^ mice.

**Supplementary Figure S5.**
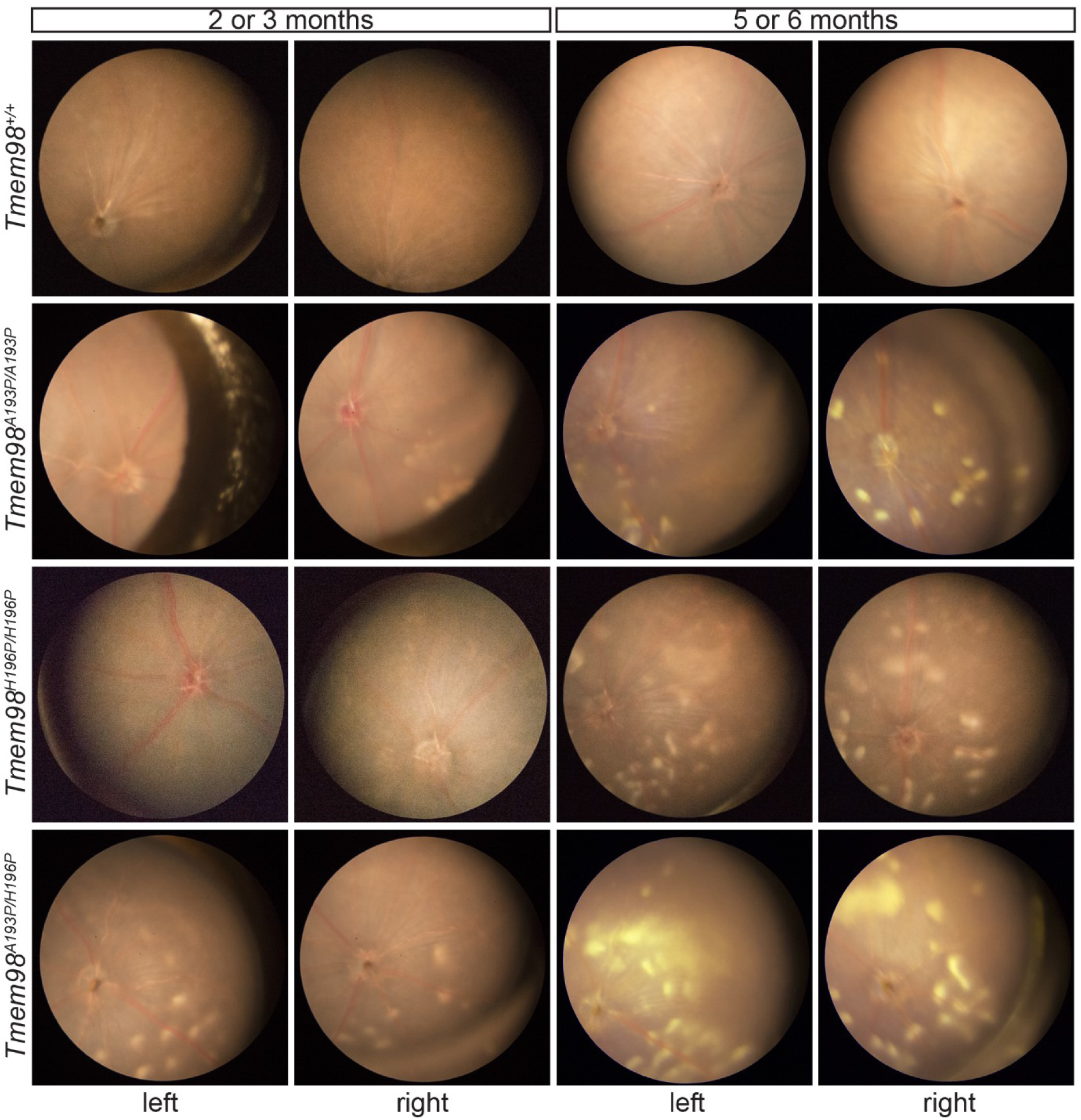
The retinal white spotting recessive phenotype in mice carrying the human nanophthalmos missense mutations is progressive. Retinal pictures of the left and right eyes of mice taken at the indicated ages. First row wild-type, second row *Tmem98*^*A193P/A193P*^, third row *Tmem98*^*H196P/H196P*^ and fourth row *Tmem98*^*A193P/H196P*^. The wild-type eyes have normal retinas at 5 months. On *Tmem98*^*A193P/A193P*^ retinas only a few spots are present at 3 months but there is extensive spotting at six months. The *Tmem98*^*H196P/H196P*^ retinas appear normal at 3 months but there is extensive spotting at six months. On *Tmem98*^*A193P/H196P*^ retinas a few spots are present at 2 months but there is extensive spotting at 5 months.

**Supplementary Figure S6.**
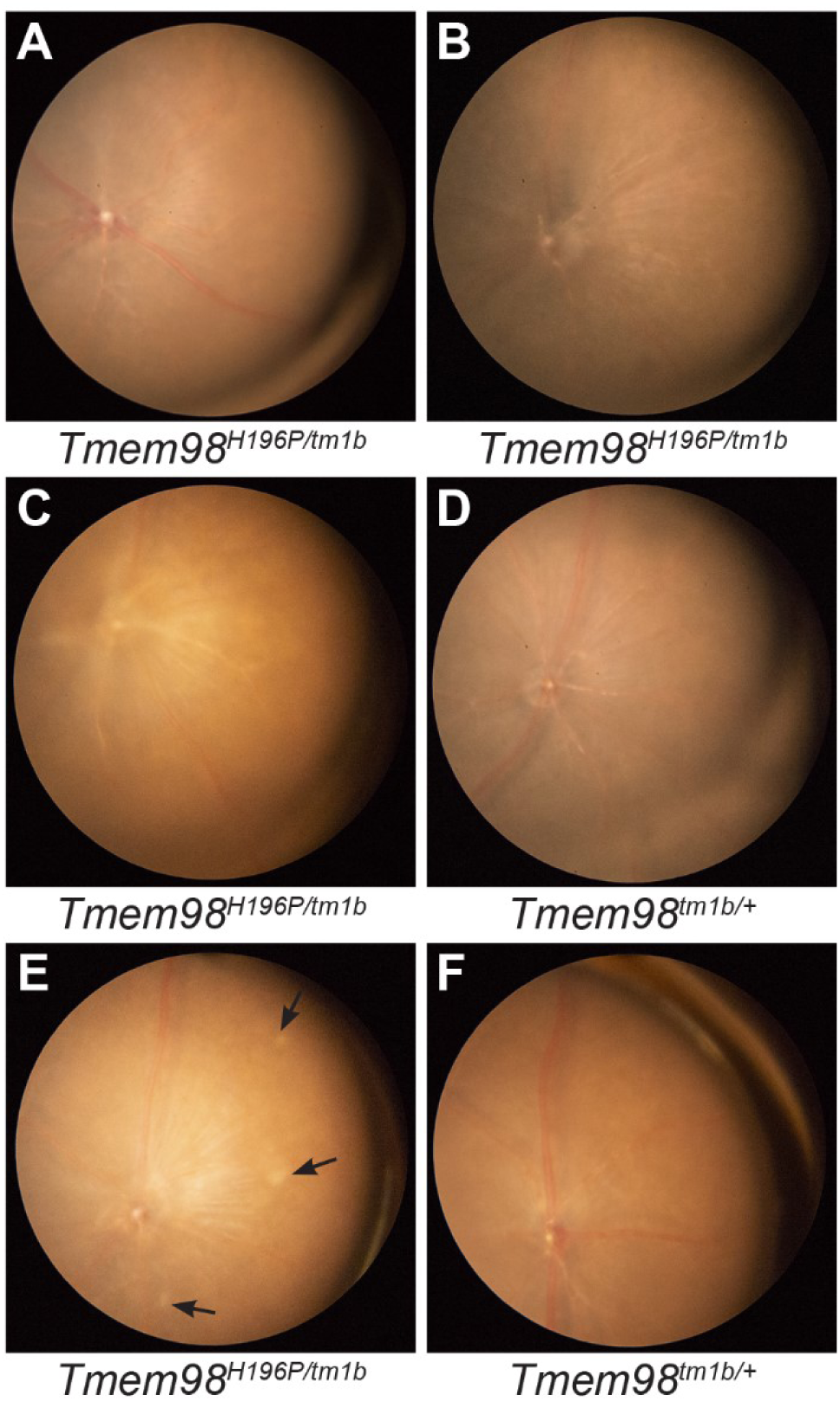
*Tmem98*^*H196P/tm1b*^ retinas rarely have retinal white spots. Retinal pictures of *Tmem98*^*H196P/tm1b*^ mice (**A**-**C** and **E**) and *Tmem98*^*tm1b/+*^ mice (**D** and **F**). The retinas shown in **E** and **F** are from littermates. *Tmem98*^*tm1b/+*^ retinas are normal. *Tmem98*^*H196P/tm1b*^ retinas are normal except for the one shown in **E** at one year of age which has three faint white spots indicated by arrows.

**Supplementary Figure S7.**
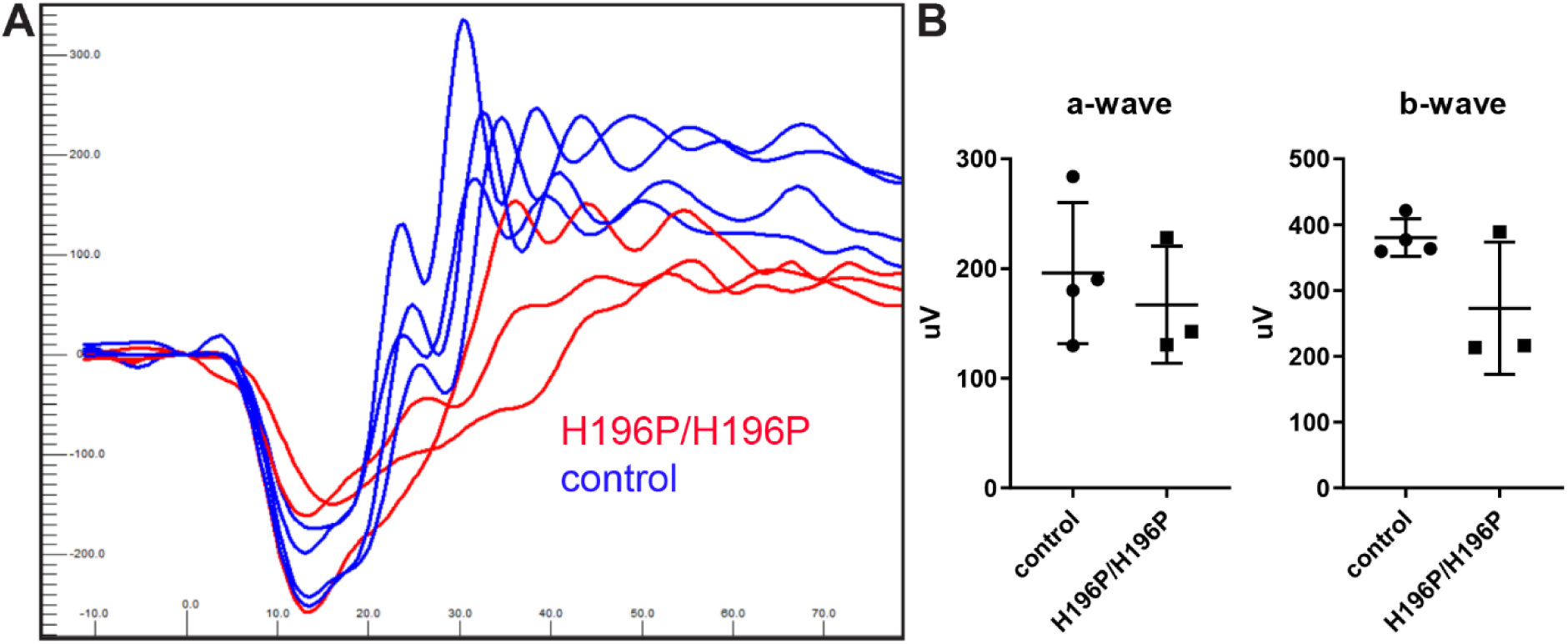
ERG response of *Tmem98*^*H196P/H196P*^ mice at 9-11 months of age is normal. Three *Tmem98*^*H196P/H196P*^ mice (two at 11 months of age and one at 9 months of age) and four 11 month old control mice (three *Tmem98*^*H196P/+*^ and one wild-type) were tested. (**A**) ERG traces of *Tmem98*^*H196P/H196P*^ mice (red lines), and control mice (blue lines). Shown are the responses at 3 cd.s/m^2^ (average of 4 flashes) for the right eye. (**B**) Comparison of a-wave amplitudes (left) and b-wave amplitudes (right), average of left and right eye for each mouse. There is no significant difference between *Tmem98*^*H196P/H196P*^ and control mice (a-wave, unpaired t test with Welch’s correction, P = 0.55 and b-wave, unpaired t test with Welch’s correction, P = 0.20).

**Supplementary Figure S8.**
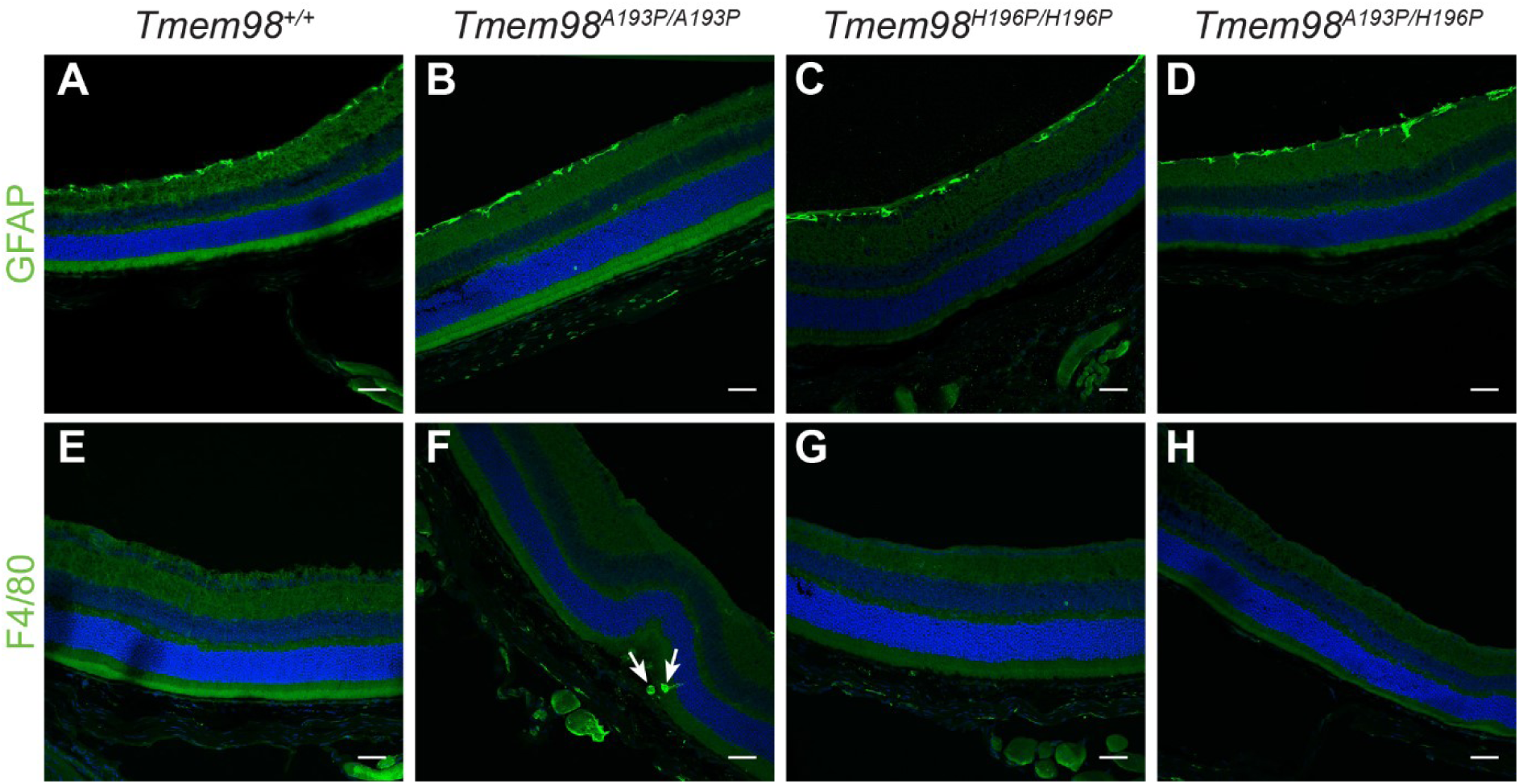
GFAP and F4/80 is normal outside the areas with retinal folds in the mutants. Immunostaining of retinal sections from wild-type mice (**A** and **E**), *Tmem98*^*A193P/A193P*^ mice (**B**, and **F**), *Tmem98*^*H196P/H196P*^ mice (**C** and **G**), *Tmem98*^*A193P/H196P*^ mice (**D** and **H**). (**A**-**D**) GFAP staining (green) is normal for all genotypes. GFAP is expressed in the ganglion cell layer where it is principally found in astrocytes. (**E**-**H**) F4/80 staining (green) was not observed outside the folds in the mutant retinas. The white arrows indicate macrophages within a fold in the outer segments for *Tmem98*^*A193P/H196P*^. (**F**) DAPI staining is shown in blue. Scale bars: 50 µm.

